# Cohesin modulates DNA replication to preserve genome integrity

**DOI:** 10.1101/2021.12.06.471507

**Authors:** Jinchun Wu, Yang Liu, Zhengrong Zhangding, Xuhao Liu, Chen Ai, Tingting Gan, Yiyang Liu, Jianhang Yin, Weiwei Zhang, Jiazhi Hu

## Abstract

Cohesin participates in loop formation by extruding DNA fibers from its ring-shaped structure. Cohesin dysfunction eliminates chromatin loops but only causes modest transcription perturbation, which cannot fully explain the frequently observed mutations of cohesin in various cancers. Here, we found that DNA replication initiates at more than one thousand extra dormant origins after acute depletion of RAD21, a core subunit of cohesin, resulting in earlier replicating timing at approximately 30% of the human genomic regions. In contrast, CTCF is dispensable for suppressing the early firing of dormant origins that are distributed away from the loop boundaries. Furthermore, greatly elevated levels of gross DNA breaks and genome-wide chromosomal translocations arise in RAD21-depleted cells, accompanied by dysregulated replication timing at dozens of hotspot genes. Thus, we conclude that cohesin coordinates DNA replication initiation to ensure proper replication timing and safeguards genome integrity.

## Introduction

The mammalian genomes are organized into different hierarchies to achieve DNA metabolism activities. The chromatin loops are fundamental units of the 3D genome and play important roles in regulating gene expression (Bonev and Cavalli, 2016; Fudenberg et al., 2016; Rao et al., 2017). Cohesin and CTCF co-anchor chromatin loops and are essential for stabilizing loop domains or topologically associating domains (TADs) (Fudenberg et al., 2016; Nora et al., 2017; Rao et al., 2017; Schwarzer et al., 2017). The ring-shaped cohesin comprises SMC1, SMC3, RAD21, and STAG1/STAG2 subunits (Gruber et al., 2003) (Figure 1A). The structural basis of cohesin in maintaining 3D genome integrity has been extensively investigated by the depiction of how cohesin translocate and extrudes DNA (Bauer et al., 2021; Davidson et al., 2019; Li et al., 2020; Shi et al., 2020).

**Figure 1.**
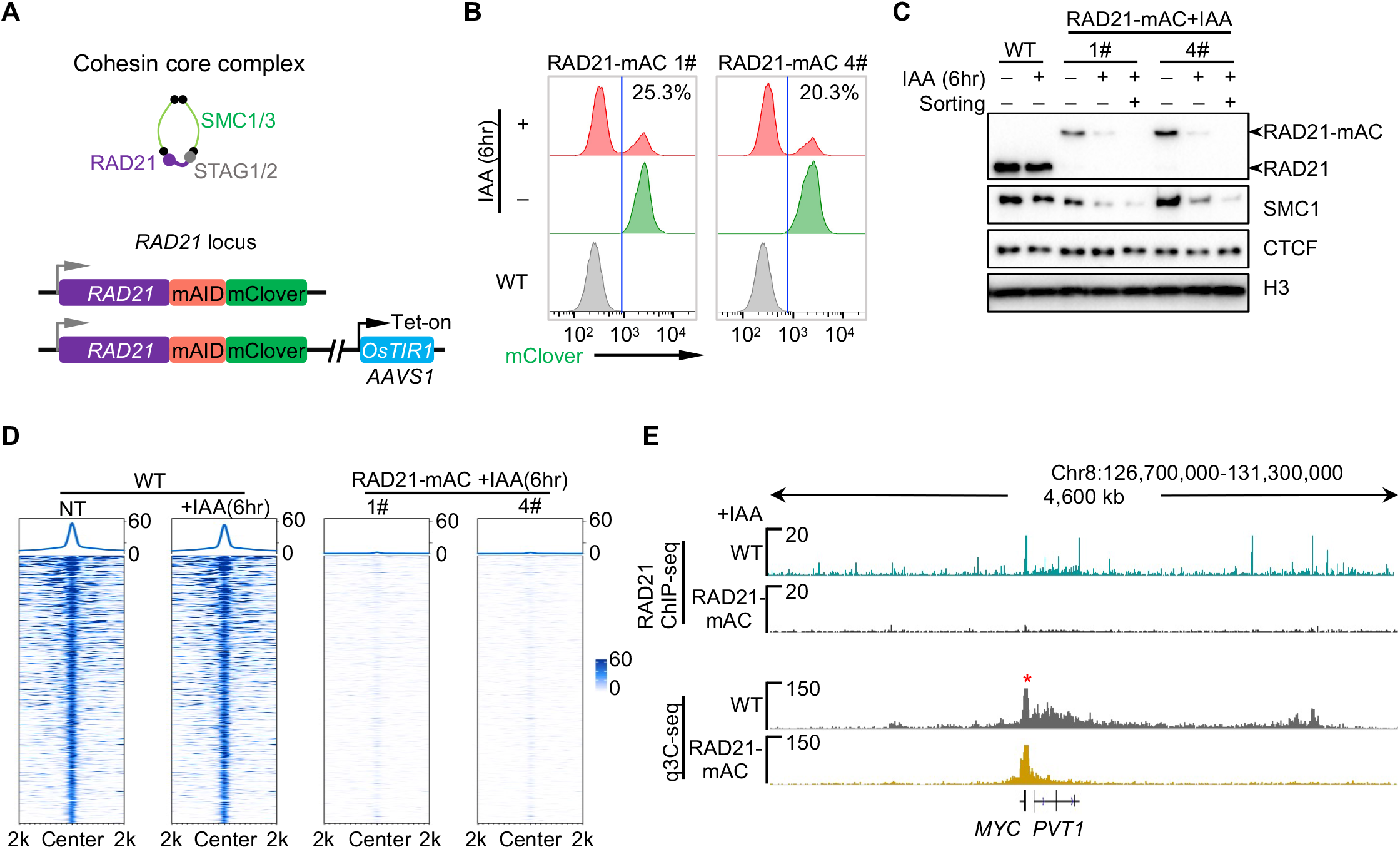
Acute depletion of RAD21 in K562 cells. A. Diagram of core subunits of cohesin (top) and the auxin-inducible degron (AID) system for rapid degradation of RAD21 (bottom). B. Indole-3-acetic acid (IAA) induces incomplete depletion of RAD21-mAC. Grey and green panels refer to the wild-type (WT) and RAD21-mAC populations, respectively, and red panels refer to RAD21-mAC clones (1# or 4#) with IAA treatment for 6 hours. mClover-positive (with percentage) and -negative cells are separated by blue lines. C. Western blotting showing the chromatin-bound indicated proteins in IAA-treated cells with or without FACS. D. RAD21 ChIP-seq signals in G1-arrested WT and RAD21-mAC cells with and without IAA treatment for 6 hours. mClover-negative RAD21-mAC cells were sorted for ChIP-seq analysis. E. Profiles of RAD21 distribution and chromatin interactions around the *c-MYC* locus in WT and RAD21-mAC cells (clone 4#) with IAA treatment for 6 hours. mClover-negative cells of RAD21-mAC were sorted for q3C-seq and ChIP-seq analysis. Red asterisk indicates the *c-MYC* locus bait for q3C-seq.

Mutations of cohesin subunits are frequently observed in various cancers including leukemia and lymphomas (Barber et al., 2008; De Koninck and Losada, 2016; Kon et al., 2013). Though cohesin is crucial for chromosome segregation, the correlation between aneuploidy and cohesin mutations in cancer cells is tenuous (De Koninck and Losada, 2016; Kon et al., 2013; Waldman, 2020). The dysfunction of cohesin leads to the elimination of loop domains and TADs, however, global transcription is mildly affected in the absence of cohesin (Rao et al., 2017). The limited ectopic gene activation induced by cohesin inactivation does not match the frequently observed cohesin mutations in cancer cells. Thus, the global contribution of cohesin dysfunction in genome integrity and tumorigenesis has yet to be fully elucidated.

DNA replication timing program is extremely robust and conserved (Marchal et al., 2019; Müller and Nieduszynski, 2012; Ryba et al., 2010), for timely and orderly duplication of the genome. Dysregulation of the replication process is the major source of genome instability and often leads to cancers (Song et al., 2018; Tomasetti et al., 2017). Cohesin has been reported to colocalize with the double hexamers of mini-chromosome maintenance complex (MCM) that is the core helicase of DNA replication machinery in the human genome (Guillou et al., 2010; Zheng et al., 2018). Moreover, abolishing the acetylation of cohesin subunit SMC3 slows replication fork progression (Terret et al., 2009), further suggesting the to-be-explored role of cohesin in DNA replication.

Employing the degron system to rapidly deplete RAD21 (Natsume et al., 2016), we explored the role of cohesin in DNA replication and genome integrity. We conclude that cohesin governs replication timing via modulating early replication initiation. And the loss of RAD21 leads to exacerbated DNA damages and chromosomal translocations within cancer-related genes in the genome, accompanied by disordered replication timing, shedding light on the mechanism for tumorigenesis of previously identified cohesin mutations.

## Results

### Acute depletion of RAD21 in K562 cells

We introduced a mini auxin-inducible degron (mAID) system to rapidly degrade the cohesin core subunit RAD21 (Natsume et al., 2016) (Figures 1A and S1A). To this end, we knocked in the doxycycline (dox)-inducible *OsTIR1* gene in the *AAVS1* locus in K562 cells and subsequentially introduced the in-frame mAID-mClover encoding sequence at the 3’ end of *RAD21* gene in both alleles (Figure 1A). Two clones 1# and 4# are identified and validated by both Sanger sequencing and western blot (Figure S1A and S1B; termed as RAD21-mAC hereafter). In the RAD21-mAC cells, CTCF and SMC1 are bound to chromatin normally (Figure S1B). In the presence of dox and indole-3-acetic acid (IAA), the RAD21-mAID-mClover protein underwent rapid degradation and more than 60% of the RAD21-mAC cells showed undetectable levels of fusion protein at one-hour post-treatment (Figure S1C). However, more than 20% or 24% of RAD21-mAC cells showed comparable expression levels of RAD21 to non-treated samples at 6- or 24-hour post-treatment, respectively (Figures 1B and S1C). To preclude the contamination of IAA-resistant cells, we isolated homogenous RAD21-depleted cells with undetectable mClover fluorescence through fluorescence-activated cell sorting (FACS) for the following analysis (Figure 1C).

To further validate the complete depletion of RAD21, we performed RAD21 chromatin immunoprecipitation-sequencing (ChIP-seq) in both wild-type (WT) and RAD21-mAC cells. Cohesin distribution in the genome was identical between IAA-treated and untreated WT cells (Figure 1D). However, RAD21 was no longer associated with its binding sites in the sorted RAD21-mAC cells treated with IAA for 6 hours (Figure 1D). Consequentially, the amount of chromatin-associated SMC1 was reduced, but CTCF remained constant on the chromatin (Figure 1C), consistent with the previous report(Rao et al., 2017). We also employed a quantitative 3C-seq (q3C-seq) to examine the impact of RAD21 loss on chromatin loops. We placed a primer anchored at the proximal upstream CTCF-binding element (CBE) of *c-MYC* and found that chromatin interaction involved in this CBE was eliminated in RAD21-depleted cells (Figure 1E). Collectively, these data indicated that RAD21 could be effectively depleted in RAD21-mAC K562 cells.

### Aberrant DNA replication initiation in RAD21-depleted cells

To determine the impact of RAD21 depletion on DNA replication, we employed previously developed nucleoside analog incorporation loci sequencing (NAIL-seq) to identify the early replication initiation zones (ERIZs) in RAD21-depleted cells (Liu et al., 2021b). Briefly, cells were synchronized at the G1 phase by CDK4/6 inhibitor for 36 hours and then treated with IAA for 6 hours before being released into G1/S transition for EdU-then-BrdU labeling (Figure S2A). The arrested RAD21-depleted cells entered the S phase normally though at a slower progression than WT cells (Figure S2B). To capture early DNA replication initiation, we collected cells at 4- and 3-hours post-release for RAD21-depleted and WT cells, respectively, with similar timing in the previous report(Liu et al., 2021b). With regards to EdU incorporation under hydroxyurea (HU) treatments, the G1-arrested WT or RAD21-depleted cells were released into HU-containing medium for 24 hours before harvest(Liu et al., 2021b) (Figure S2A). Of note, the amounts of origin recognition complex (ORC) and MCM remained constant on the chromatin before release (Figure S2C).

We identified 2,122 ERIZs in IAA-treated WT cells, similar to our previous report(Liu et al., 2021b). While 3,593 and 3,475 ERIZs were identified in RAD21-mAC clones 1# and 4#, respectively, with a very high correlation (Figures 2A, 2B, and S2D). Strikingly, 1,209 and 1,095 new ERIZs were identified in RAD21-depleted 1# and 4# cells, respectively, which occupied approximately 30% of the total ERIZs in each RAD21-depleted clone (Figure 2B-D). By comparing the replication signal intensity of shared ERIZs between WT and RAD21-depleted cells, we identified the “increased” (639 in 1# and 630 in 4#), “equal” (1,555 in 1# and 1,544 in 4#), and “decreased” (190 in 1# and 206 in 4#) ERIZs that showed up-regulated, equivalent, or down-regulated early replication signals in RAD21-depleted cells in comparison to WT cells (Figures 2A-D). Besides, 202 or 203 “disappeared” ERIZs occurred in WT but not in RAD21-depleted cells. However, these disappeared ERIZs were relatively weak in WT cells and the total replication signal intensity was only slightly decreased in RAD21-depleted cells (Figure S2E), so we combined them into “decreased” ERIZs for further analysis (Figure 2B). In total, only less than half of the ERIZs exhibited comparative intensity of replication signals after RAD21 depletion (Figure 2C).

**Figure 2.**
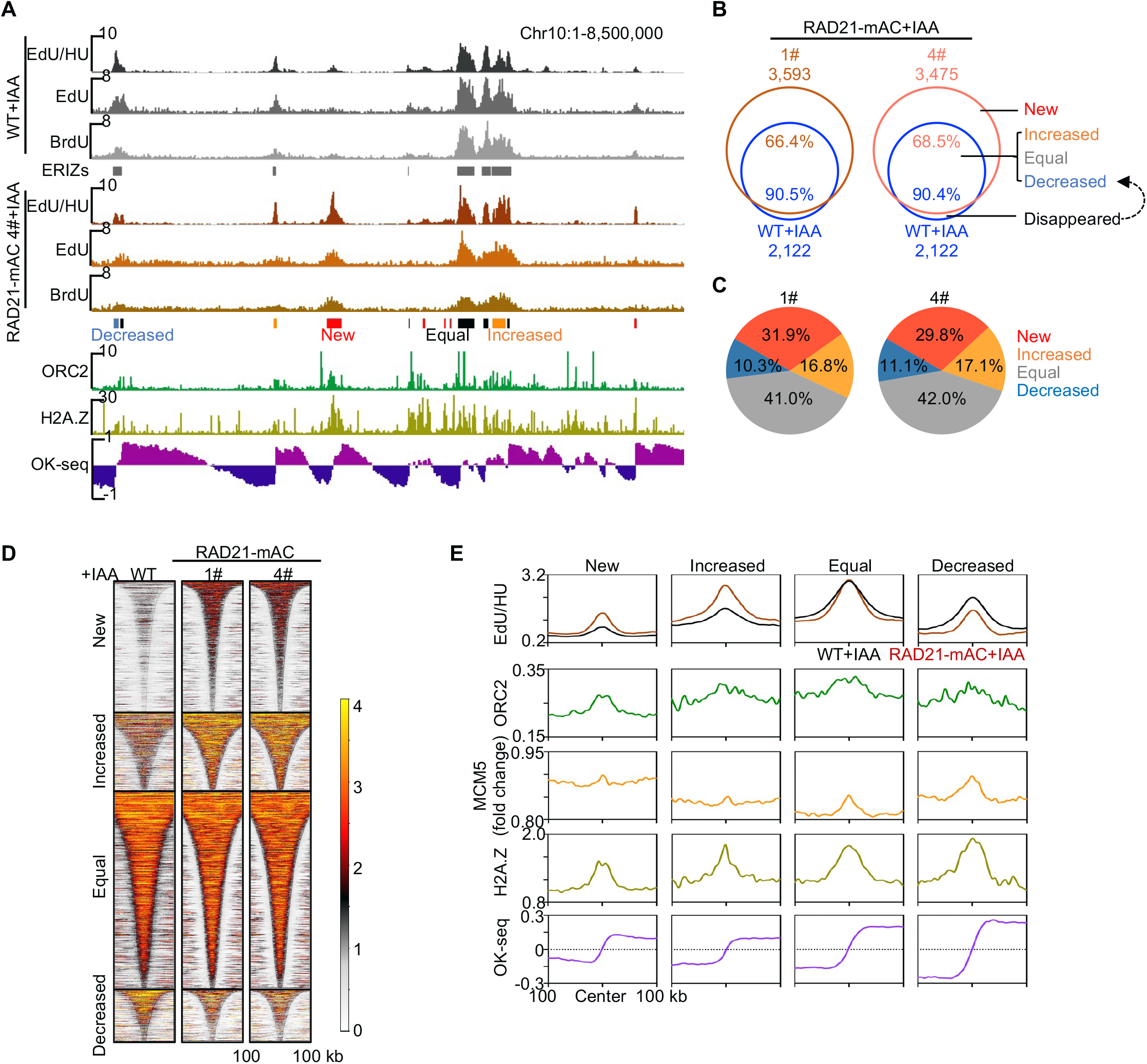
RAD21 regulates early replication initiation in K562 cells. A. Representative profiles of early replication signals and identified ERIZs in WT and RAD21-depleted cells (clone 4#). The ChIP-seq signals of ORC2, H2A.Z, and OK-seq of WT cells from previous reports (Miotto et al., 2016; Wu et al., 2018) were on the bottom. B. Venn diagram showing the overlap of ERIZs between WT and RAD21-depleted cells from indicated clones. The black curved arrow represented that the “disappeared” ERIZs were merged into “decreased” ERIZs for further analysis. C. Percentages of each class of ERIZs after RAD21 depletion in clone 1# and 4#, respectively. D. Heatmaps showing the distribution of early replication signals from IAA-treated WT and RAD21-mAC cells at the ERIZs of RAD21-mAC 1#. The ERIZs were ranked by width and centered at the midpoint. E. Distributions of EdU/HU, ORC2, H2A.Z, and OK-seq at the 4 classes of ERIZs identified from RAD21-mAC 1#. ERIZs are aligned at the mid-point. ORC2, MCM5, H2A.Z, and OK-seq were from WT cells. And ORC2, H2A.Z, and OK-seq were from re-analysis of previous data (Miotto et al., 2016; Wu et al., 2018).

We next examined the chromatin features of the new ERIZs identified in RAD21-depleted cells. We found that the replication initiation-associated factors, ORC, MCM, and H2A.Z (Long et al., 2020) were enriched at the new ERIZs, similar to other classes of ERIZs in RAD21-depleted cells (Figure 1E). In addition, transcription was also devoid in all four classes of identified ERIZs including the new ones, in line with previous reports that replication initiation occurred in non-transcribed regions (Liu et al., 2021b; Wang et al., 2021) (Figure S2F). A sharp switch of OK-seq signals was also detected right at the center of the new ERIZs (Figure 1E). Collectively, the new ERIZs in RAD21-depleted cells show typical features of DNA replication origins, suggesting that they may originate from temporal dysregulation of origin firing after RAD21 depletion, but not occur *de novo*.

### Perturbed DNA replication timing in RAD21-depleted cells

To determine whether the new ERIZs result from dysregulated origin firing, we performed replication timing analysis for RAD21-depleted cells. To this end, we labeled the growing WT or RAD21-depleted cells with EdU for 15 minutes and then collected the cells into 6 fractions by DNA content as previously described (Brison et al., 2019) (Figure S3A). We used the EdU labeling method in NAIL-seq to construct sequencing libraries and adapted the Repli-seq pipeline for replication timing computation (Brison et al., 2019; Liu et al., 2021b). The new ERIZs induced advanced replication of the neighbor regions in RAD21-depleted cells, resulting in relatively earlier replication timing than the same regions in WT cells (Figures 3A, 3B, and S3B). The two replicates of WT cells showed resemble profiles of replication timing and we used their correlation map to determine the 95% quantile interval for the assessment of unchanged regions (Figure 3C, left panel). Approximately 10.2% and 30.1% of the genomic regions showed significantly earlier replication timing in RAD21-depleted cells treated with IAA for 6- or 24-hours, respectively (Figures 3C right panel, S3B and S3C). In the RAD21-depleted 4# cells at 24-hours post-IAA-treatment, 527 out of 1,095 new ERIZs induced advanced replication in the adjacent regions, while 168 increased and 103 equal ERIZs led to earlier replication timing (Figure 3D). Of note, 42 decreased ERIZs exhibited earlier replication timing, possibly because they were close to other classes of ERIZs (Figures 3D and S3E). Thus, cohesin dysfunction leads to aberrant replication timing in K562 cells.

**Figure 3.**
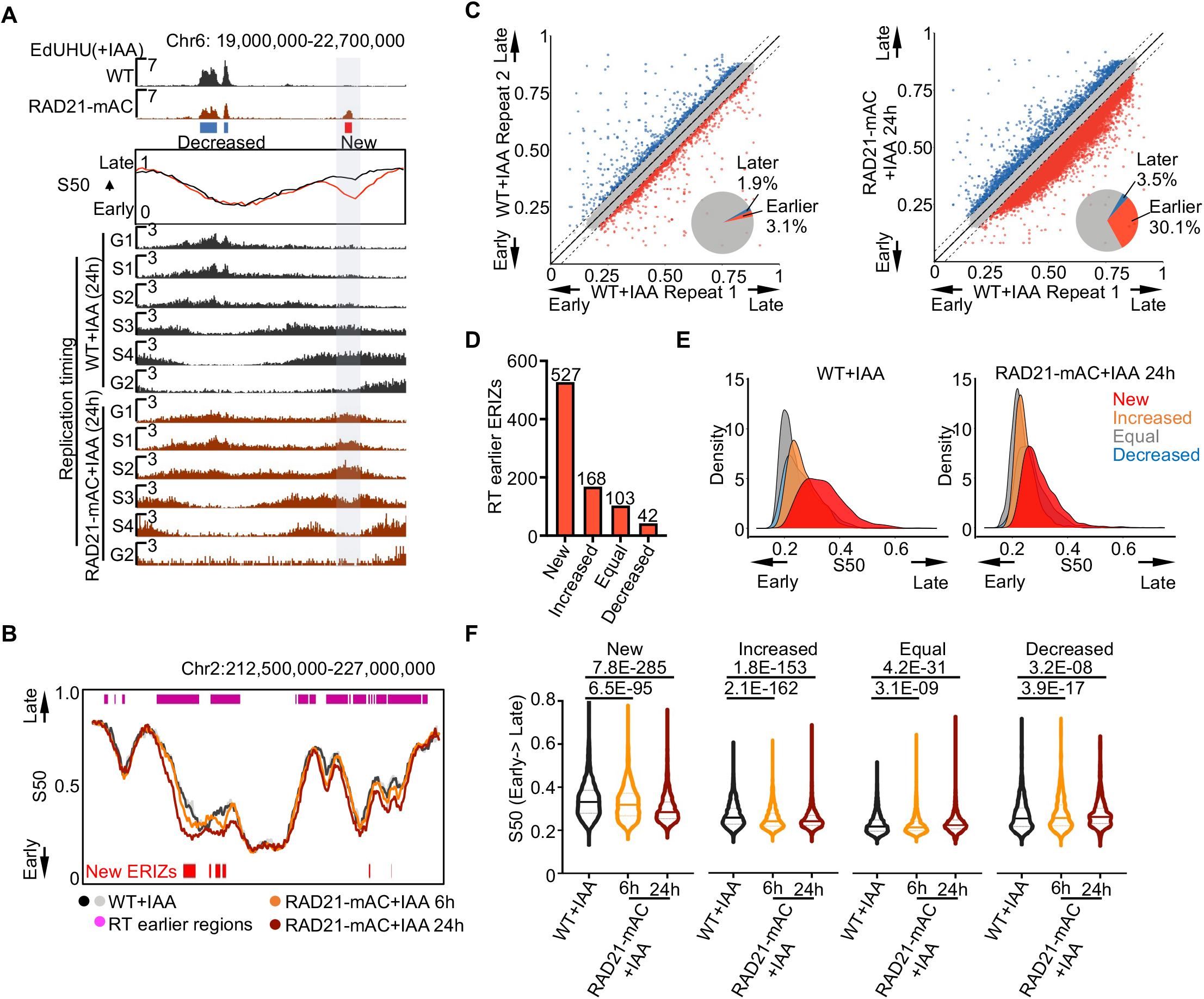
RAD21 depletion reprograms replication timing in K562 cells. A. Representative example of a “new” ERIZ nested in an earlier replication timing region. The profiles of replication timing were represented by S50. The black curve is WT+IAA for 24h and the dark red line is RAD21-mAC 4#+IAA for 24h. Bin size=50 kb. Replication signals were placed on the bottom. Shadow in grey indicates the position of new ERIZ. B. Profiles showing the replication timing in a 13.5-Mb region. Replication timing profiles were from WT (black or grey lines, IAA for 24 h) and RAD21-mAC 4# (orange or dark red line, IAA for 6h or 24 h, respectively) cells. Red strips mark the “new” ERIZs identified in Figure1 d-f; magenta strips refer to the replication timing significantly advanced regions identified in Figure 3C. C. Left: Scatter plot of S50 value between two WT+IAA replicates. Right: Scatter plot of S50 value between WT replicate 1 and RAD21-mAC cells (clone 4#) with IAA treatment for 24 hours. Each dot represents a 50-kb genomic region. Grey dots within the two dashed lines represent unchanged regions in WT+IAA cells. Blue or red dots mark bins with a later or earlier replication timing in the WT+IAA replicate 2 (left) or RAD21-mAC+IAA (right), respectively. The pie charts summarize the percentages of bins with a later (blue) or earlier (red) replication timing in each panel. D. The numbers of ERIZs (from RAD21-mAC 4#) locate in earlier timing regions identified in Figure 3C. The percentages of earlier ERIZs in “new”, “increased”, “equal”, and “decreased” ERIZs are 48.1%, 26.7%, 6.7%, and 10.3%, respectively. E. Density plots showing the distribution of S50 values of four classes of ERIZs identified from RAD21-depleted cells (clone 4#) in the WT (left) or RAD21-mAC (right) replication timelines, respectively. F. The distribution of S50 values of four classes of ERIZs from WT or RAD21-depleted cells (clone 4#). The Wilcoxon matched-pairs signed-rank test was applied for statistics and the *p*-value is marked on the top.

To determine the original replication timing of the ERIZs identified from RAD21-depleted cells, we plotted these ERIZs on the replication timeline of RAD21-proficient cells. The initiation of equal ERIZs occurred at similar time points between WT and RAD21-depleted cells while decreased ERIZs showed a minor delay in the RAD21-depleted cells with IAA treatment for 24 hours (Figures 3E and S3F). In contrast, the genomic regions containing the increased ERIZs replicated earlier and the new ERIZs-containing regions showed an even more significant earlier replication timing after RAD21 degradation (Figures 2E, 2F, and S3F). These data imply that the new ERIZs in RAD21-depleted cells arise from the earlier firing of substantial origins that are originally dormant at the early replication initiation stage.

### Redistributed replication initiation within loop domains in RAD21-depleted cells

Given that both cohesin and CTCF are required for the formation of loop domains (Figure 4A), we next sought to investigate whether CTCF is also involved in the advanced firing of identified new ERIZs. To this end, we constructed a CTCF-mAID-mClover (CTCF-mAC, clone 5C9) K562 cell line. Similar to the RAD21-mAC cell lines, the 3’ end of *CTCF* was tagged with an in-frame mAID-mClover encoding sequence on both alleles (Figures S4A, B). We isolated the CTCF-depleted cells by FACS to remove IAA-resistant cells and performed NAIL-seq EdU/HU method as for RAD21-mAC cells (Figure S2A and S4B). The overall distribution profile of replication signals at identified ERIZs in CTCF-depleted cells was similar to that in WT cells except with a lower intensity (Figures 4B-D, and S4C). Specifically, CTCF-depleted cells exhibited very low intensity of replication signals at the new ERIZs as WT cells (Figure 3C), suggesting that CTCF is dispensable for suppressing the firing of dormant origins.

**Figure 4.**
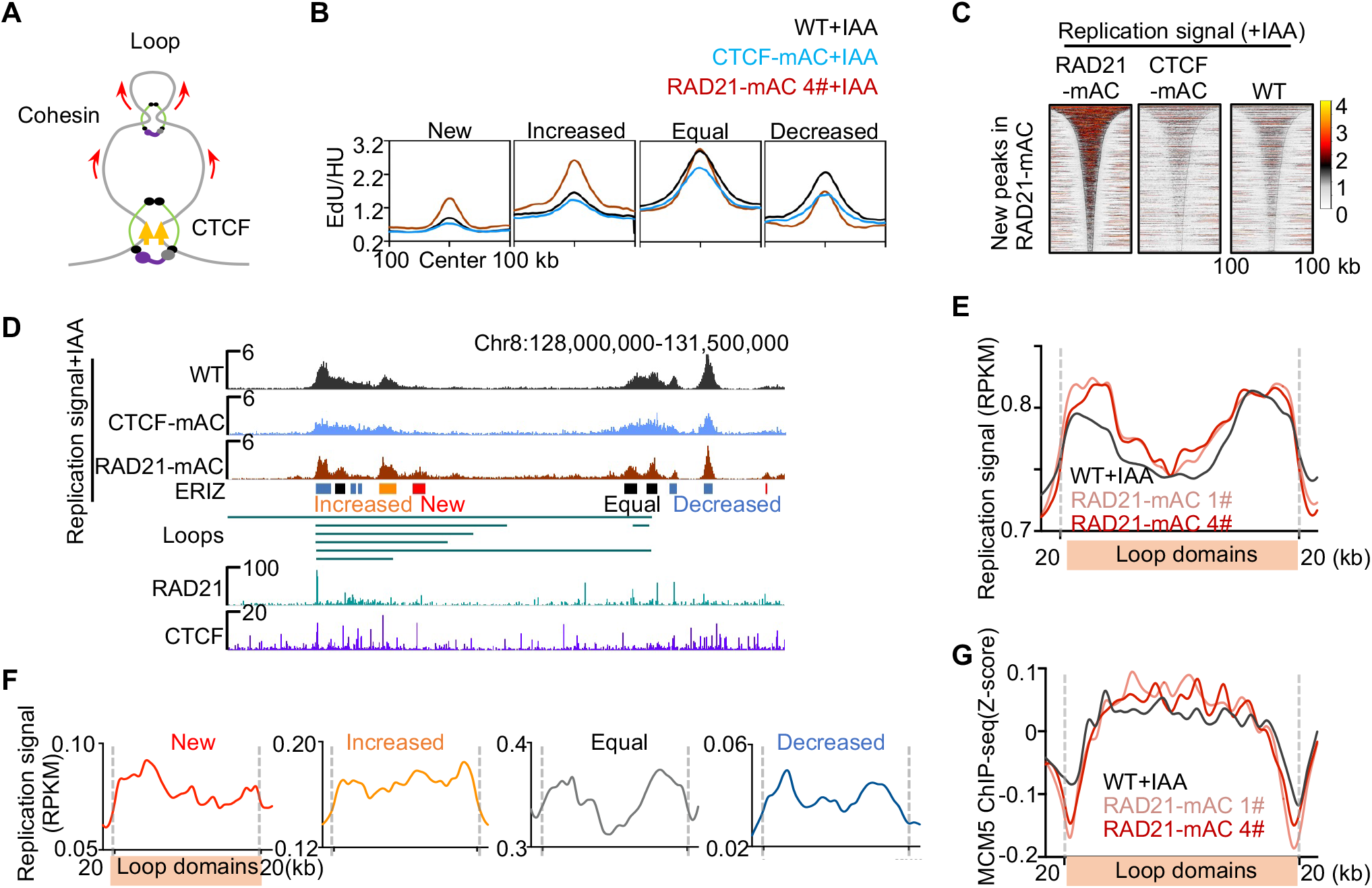
Distribution of early replication signals within loop domains. A. Diagram of Cohesin-mediated loop extrusion model. Yellow and red arrows refer to the CTCF protein and the movement direction of DNA, respectively. B. Distribution of early replication signals of IAA-treated WT, RAD21-mAC, and CTCF-mAC cells in the centered ERIZs identified in the RAD21-depleted cells (clone 1#). C. Heatmaps showing the distribution of early replication signals from IAA-treated RAD21-mAC, CTCF-mAC, and WT samples at the new ERIZs identified from RAD21-mAC (clone 1#). Legends are depicted as described in Figure 2D. D. Example of early replication signals in a represented loop domain after RAD21 or CTCF depletion. The ChIP-seq signals of RAD21 and CTCF are showed on the bottom. E. Distribution of early replication signals in the presence (black) or absence (pink or red) of RAD21 within loop domains in K562 cells. Each loop domain was divided into 250 bins and aligned at loop boundaries, with 20-kb regions upstream and downstream. F. Distribution of early replication signals in the four classes of ERIZs after RAD21 depletion within the loop domains in K562 cells. Legends are depicted as described in e. G. Distribution of MCM5 signals in the presence (black) or absence (pink or red) of RAD21 within loop domains in K562 cells. The fold-change of MCM5 is defined as the ratio of ChIP-ed over input signals and normalized to z-scores.

Cohesin functions as molecular motors to progressively extrude DNA to convergent CTCF that is resident at the boundaries of loop domains (Davidson et al., 2019) (Figure 4A), therefore we examined the distribution of early replication signals in chromatin loop domains (Rao et al., 2014). In WT cells, early replication tended to occur at the regions close to loop boundaries (Figure 4E). Though the overall early replication profile of cohesin-depleted cells was similar to that of the WT cells, more replication signals accumulated in the regions away from the loop boundaries (Figure 4E). In contrast, the replication signals within non-loop domains were highly similar between WT and RAD21-depleted cells (Figure S4D). With regards to the distribution of four classes of ERIZs in loop domains, much more replication signals from new and increased ERIZs occurred in the middle of loop domains (Figure 4F). The median distance to loop boundaries was up to 40.8 kb and 32.0 kb for new and increased ERIZs, respectively, in RAD21-depleted cells, in comparison to 26.5 kb for all ERIZs of WT cells (Figure S4E). We also performed MCM5 ChIP-seq analysis for G1-arrested cells because origin licensing occurs in the G1 phase (O’Donnell et al., 2013). Similar to the early replication signals, MCM displayed a higher accumulation in the middle of loop domains in cohesin-depleted cells than in WT cells (Figure 4G). Collectively, these data suggest that loop extruding by cohesin within the entire loop domains possibly suppresses aberrant origin firing away from loop boundaries.

### Increased chromosomal translocations in RAD21-depleted cells

To determine the impact of cohesin dysfunction on genome stability, we employed a previously developed primer-extension-mediated sequencing (PEM-seq) assay to profile and quantify the DNA double-stranded breaks (DSBs) in RAD21-depleted cells (Liu et al., 2021a; Yin et al., 2019; Zhang et al., 2021). We used CRISPR-Cas9 to generate bait DSBs at the *c-MYC* or *TP53* loci and captured the genome-wide DSBs that formed chromosomal translocations with the bait DSBs (Figure 5A). Dramatically increased chromosomal translocations were detected in RAD21-depleted cells, especially at the bait site-containing chromosome, and the total translocation levels were 3 to 5 fold of that in IAA-treated WT cells (Figures 5B, 5C, and S5A). We then identified genome-wide enriched translocation clusters in RAD21-depleted cells against WT cells with a significance of FDR<0.01. 196 and 204 translocation clusters were identified at the two bait sites, which harbored 30 to 46 hotspot genes for the two RAD21-mAC clones (Figures 5D, S5B and S5C). More than 45% of these hotspot genes were involved in translocations of both bait sites in RAD21-depleted cells (Table S1). Moreover, more than one-third of these hotspot genes were reported to be mutated in cancers or other diseases, including *KDM6A*, *SMYD3*, *RUNX1*, *MAPK1*, and *BCR* (Figures 5E, S5D, S5E, and Table S1).

**Figure 5.**
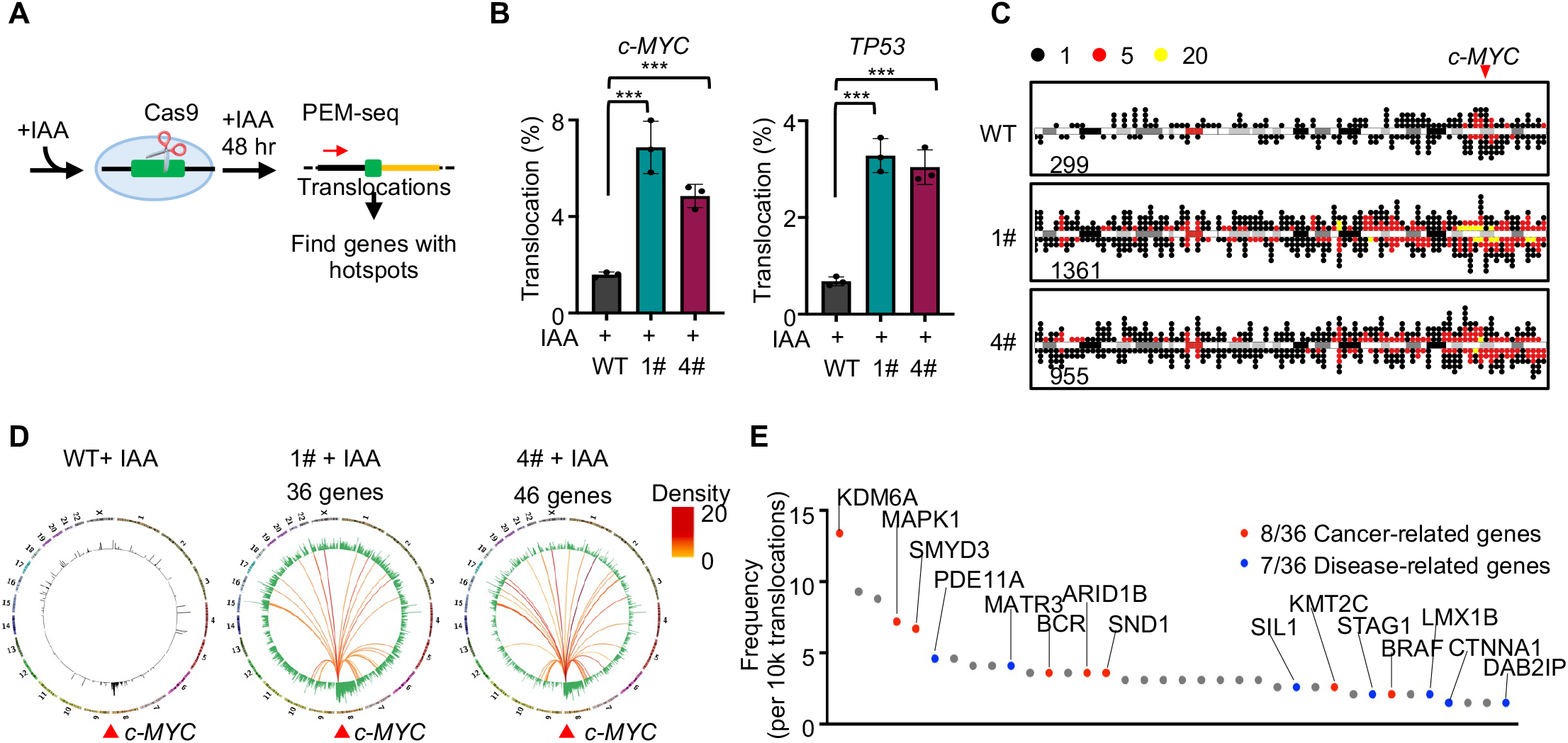
RAD21 depletion leads to increased chromosomal translocations. A. The strategy for mapping DSBs and genome-wide translocations in RAD21-depleted cells. Cells were sorted by FACS for PEM-seq analysis. B. Frequency of translocations with or without RAD21 depletion. Mean±SD from three repeats; *t*-test, \*\*\**p*<0.001. The bait sites are indicated on the top. C. Dot plots exhibiting the distribution of translocation junctions on Chromosome 8 in IAA-treated cells. The number of translocations were normalized to 100,000 editing events from each sample. The red arrow indicates the position of bait site at the *c-MYC* locus. Numbers of junctions are indicated at each plot. Legend for dot colors is indicated on the top. D. Circos plots showing the genome-wide translocations and translocation hotspot genes cloned from *c-MYC* locus. The outer circle shows each indicated chromosome of human genome. The inner circle shows the number of translocations within each 1-Mb bin with a log scale in IAA-treated WT (black) and RAD21-depleted (green) cells. Bins with less than 3 translocation junctions were considered as under background levels and were not shown. Red arrows indicate the bait site of *c-MYC*. Colored lines link the bait site and translocation hotspot genes after RAD21-loss and refer to the translocation frequency of each hotspot gene per 10,000 translocations, indicated by the color legend. E. Translocation frequencies of RAD21-depletion-induced hotspot genes in RAD21-mAC 1# captured by *c-MYC* DSBs. Cancer-related (red) and disease-related (blue) genes are annotated by The Human Protein Atlas.

We also employed precision nuclear run-on sequencing (PRO-seq) to examine the transcription profile of identified hotspot genes (Jiang et al., 2020; Sigova et al., 2015). The vast majority of these hotspot genes showed similar transcription levels in the presence or absence of RAD21 (Figure S6A), indicating that transcription is not involved in inducing DSBs at these hotspot genes. Then we examined the change of replication timing of these hotspot genes. Approximately 34.4% of total genes fell in regions with altered replication timing, while the percentages of the identified hotspots genes were increased to more than 55% at both bait sites (Figure 6A), indicating the involvement of altered replication timing in DNA breaks at these hotspot genes. Specifically, more hotspot genes exhibited earlier rather than later replication timing, exemplified by *KDM6A* and *STAG1* captured by *c-MYC* DSBs in addition to *CCNY* and *RUNX1* captured by *TP53* DSBs (Figure 6B-D).

**Figure 6.**
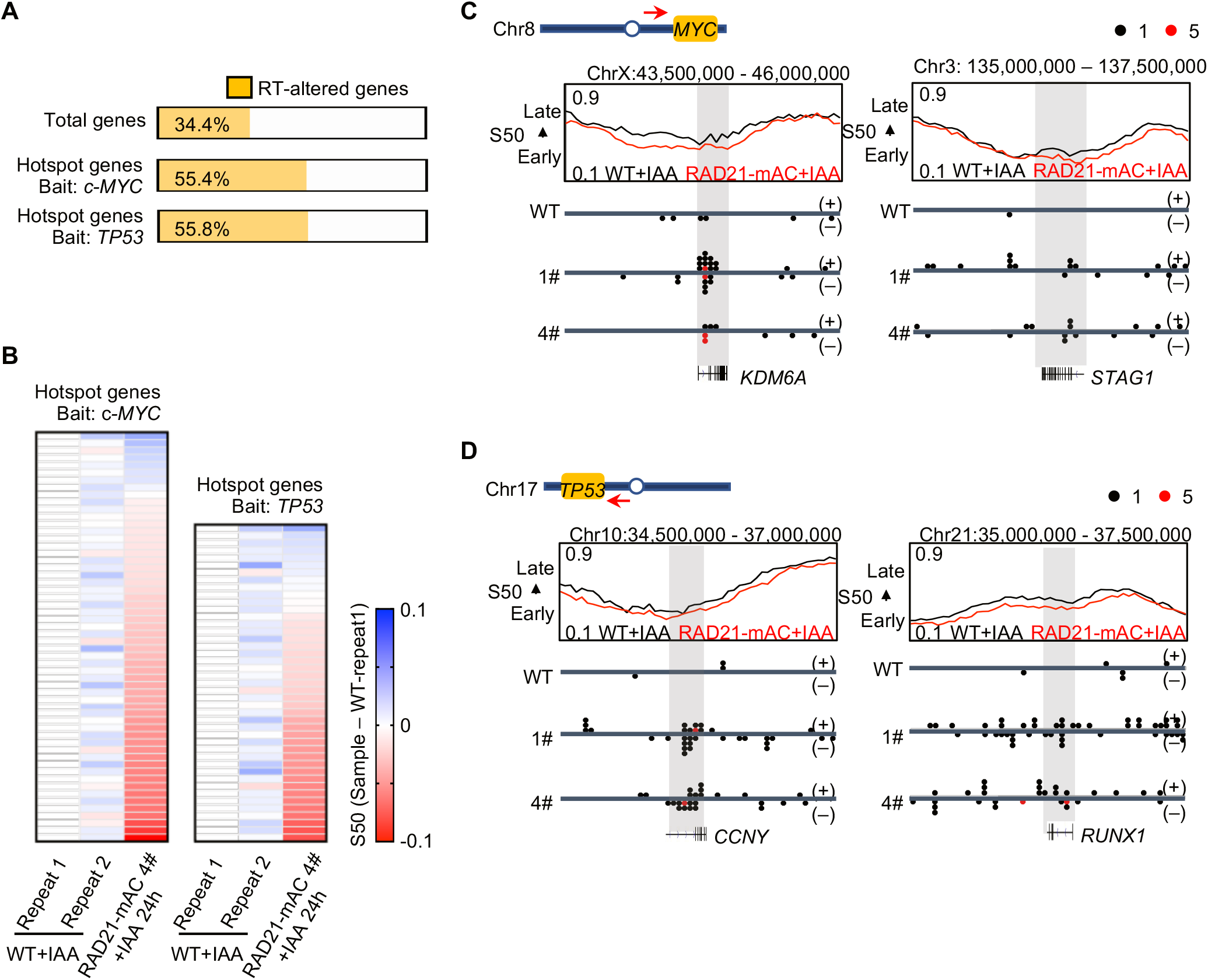
Translocation hotspot genes tend to locate in replication timing-altered regions. A. The proportion of the total annotated genes (*GENCODE*, Top) or translocation hotspot genes (Middle and Bottom) containing replication timing-altered regions. The hotspot gene list contained the hotspots genes identified from both RAD21-mAC 1# and 4# clones. RT, replication timing. B. Heatmaps showing the divergences of S50 between RAD21-depleted and IAA-treated WT cells in each translocation hotspot gene. Each row indicates a hotspot gene. The S50 for indicated gene of WT replicate 1 was defined as 0 and the S50 from other samples were normalized to WT replicate 1 (direct subtraction). The colors represent the mean value of S50 in the gene region. Red or blue refers to relatively earlier or later replication timing, respectively, in comparison to WT replicate 1. C. Examples of hotspot gene regions captured by *c-MYC* (top) and *TP53* (bottom). The profile of replication timing in indicated regions is shown on the top. The dot plots present translocation junctions identified from indicated locus. The red arrows indicate the primer direction of PEM-seq. The black and red lines mark the S50 in IAA-treated WT and RAD21-mAC cells, respectively. The grey shadows mark the hotspot genes.

## Discussion

Though the degron system enables the rapid degradation of target protein, the depletion efficiency varies for different proteins. In our system, more than 20% of RAD21-mAC K562 cells are resistant to IAA treatment, in line with a previous report in HCT116 cells (Oldach and Nieduszynski, 2019) (Figure 1B and S1C). Given that RAD21-proficient cells enter the S phase more efficiently than RAD21-deficient cells (Figure S2C), the impact of these RAD21-retained cells on ERIZ identification and replication timing profiling can be further exacerbated. Therefore, we employed FACS to isolate homogenous RAD21-deficient cells for all the analyses to rule out the contamination of both RAD21-retained cells and dead cells.

Dormant origins are fired in the early S phase after RAD21 depletion, which leads to perturbed replication timing. We identified 50% more ERIZs in RAD21-depleted cells in comparison to WT cells, consistent with the previous report (Cremer et al., 2020). The newly identified ERIZs in RAD21-depleted cells are distributed evenly in the loop domains, different from the ERIZs of WT cells that are enriched in the regions close to the loop boundaries (Figure 4E, 4F). Moreover, cohesin suppresses the firing of dormant origins, which is independent of CTCF (Figure 4C). Therefore, the CTCF-dispensable loop extruding function of cohesin is important for ensuring proper firing of early DNA origins. In this context, cohesin may lead to redistribution of MCM during loop extrusion to promote DNA replication initiation at the regions close to loop boundaries in WT cells (Figure S6B). While in the absence of cohesin, MCM may accumulate at any region within the loop domains, which results in more DNA replication initiation in the middle of loop domains (Figures 4E-G, and S6D). With this regard, we detected a coincident redistribution of MCM and early replication signals in the middle of loop domains after cohesin dysfunction (Figure 4E and 4G). Moreover, MCM double hexamer colocalizes with cohesin in Hela cells, which can be detected during MCM redistribution by cohesin (Guillou et al., 2010; Zheng et al., 2018).

Similarly, cohesin has been proposed to redistribute the RAG complex during V(D)J recombination in developing lymphocytes (Dai et al., 2021; Hu et al., 2015; Zhang et al., 2019). Nevertheless, how cohesin interplays with MCM remain to be explored.

Excess DNA replication initiation induced by RAD21 depletion may pose great threats to genome integrity. Replication initiation *per se* may occasionally cause DSBs because accumulating evidence showed that early replication is colocalized with double-stranded DNA breaks and DNA repair factors in mouse B cells (Barlow et al., 2013; Tubbs et al., 2018). Moreover, excess replication forks consume more replication factors than normal, and the exhausted replication limiting factors lead to severe replication stresses (Toledo et al., 2013; Zeman and Cimprich, 2014). In addition, a collision between replication forks and transcription as well as excess replication terminations may be also involved in the genome instability after cohesin dysfunction (Hamperl et al., 2017; Liu et al., 2021b; Wang et al., 2021). In this context, the alteration of replication timing program is prevalent in some cancers and associated with common fragile sites (Brison et al., 2019; Ryba et al., 2012). Therefore, cohesin dysfunction induces greatly elevated levels of DNA breaks and chromosomal translocations. At the tested two loci, the genome-wide translocation levels are increased to 3 to 5 fold in RAD21-depleted K562 cells (Figure 5B), even higher than the 2-fold change detected in the ATM-deficient B cells (Hu et al., 2014). Of note, ATM is the mater kinase for DSB repair and ATM mutations frequently lead to lymphomas and leukemia in both humans and mice (Alt et al., 2013; Gostissa et al., 2011).

## STAR Methods

### Cell culture, mAID-mClover introducing, and cell cycle synchronization

The authenticated K562 cells (3111C0001CCC000039, National Infrastructure of Cell line resource, China) were cultured in the RPMI1640 with 15% FBS as described (Liu et al., 2021b). The *OsTIR1* gene controlled by the Tet-On promoter was integrated into the *AAVS1* locus by CRISPR-Cas9-mediated homology-directed repair (HDR). Clones were validated for the integration at the target site by both PCR and western blotting. A validated clone with a higher expression level of OsTIR1 than other clones was subjected to the subsequential gene manipulation.

For the tagging of either RAD21 or CTCF, the selected clone expressing OsTIR1 was transfected with plasmids containing Cas9-single guide RNA and an in-frame mAID-mClover tag, flanked by homology arms identical to RAD21 or CTCF, respectively, by nucleofector (Lonza, 4D-Nucleofector X). Cells were subcloned and candidate clones were subjected to further validation by PCR, sanger sequencing, flow cytometer analysis, and western blot as presented. Finally, two clones (RAD21-mAC 1# and 4#) for RAD21 and one clone (CTCF-mAC 5C9) for CTCF were validated and used for further analysis.

The WT, RAD21-mAC, and CTCF-mAC are synchronized as previously reported(Liu et al., 2021b). For G1 arrest, cells were incubated with 5 μM Palbociclib (Selleckchem, S1116) for 36 hours. To be released to the G1/S transition, G1-arrested cells with or without RAD21 were released into a fresh medium for 4 or 3 hours, respectively. For HU treatment, G1-arrested cells were released into a fresh medium supplied with 10 mM HU for 24 hours. For detailed treatments of each assay including ChIP-seq, q3C-seq, NAIL-seq, Repli-seq, PEM-seq, and PRO-seq, please refer to the following sections. All the sequencing data were aligned to human genome hg19 for further analysis.

### ChIP-seq of RAD21, CTCF, and MCM5 in the G1 phase

For G1 arrest, both WT and RAD21-mAC (1# and 4#) cells were cultured with palbociclib (Selleckchem, S1116) for 36 hours. Cells with no treatment were set as control. To induce the acute depletion of RAD21, 2ug/ml dox (Sigma, D9891) and 500uM IAA (Sigma, I5148) was included in the medium at 24 hours and 6 hours before harvest, respectively, and cells were then subjected to FACS purification for the mClover-negative cells indicating the ablation of RAD21. Only G1-arrested WT cells were applied to CTCF ChIP-seq analysis.

Collected cells were fixed by 1% formaldehyde (Sigma, F1635, fresh-made) for 10 min at 25°C and quenched by 125 mM glycine (VWR-amerasco 0167) for 5 min. Thoroughly washed by PBS, cells were lysed on ice for 15 min with ice-cold NP40 lysis buffer (10 mM Tris-HCl, pH 7.5; 150 mM NaCl; 0.05% NP40) supplied with protease inhibitors (PIs) (Bimake, B14012). The nuclei fraction was isolated, washed by PBS/1 mM EDTA, and then lysed with glycerol buffer (20 mM Tris-HCl, pH 8.0; 75 mM NaCl; 0.5 mM EDTA; 0.85 mM DTT; 50% glycerol) and nuclei lysis buffer (10 mM HEPES, pH 7.6; 1 mM DTT; 7.5 mM MgCl2; 0.2 mM EDTA; 0.3 M NaCl; 1 M urea; 1% NP40). The chromatin was pelleted by centrifuge, washed twice by PBS/1 mM EDTA, and eventually suspended in sonication buffer (20 mM Tris-HCl, pH 8.0; 150 mM NaCl; 2 mM EDTA; 0.1% SDS; 1% Triton X-100; 4 mM CaCl2) with PIs. Chromatin fractions were then treated with 50 U MNase (NEB, M2047S) for 10-12 min at 37°C, and the digestion was quenched by adding 5 mM EGTA and 5 mM EDTA. After that, pre-digested chromatin was sonicated by a Biorupter (Energy: High; On: 30 s; Off: 60 s; 15 cycles; 4°C). 30 μL of soluble chromatin fraction was kept as input and the rest was subjected to ChIP analysis.

For ChIP-seq, an antibody against RAD21 (1:50; Abcam, ab992), CTCF (1:100; Millipore 07-729) or MCM5 (1:50; Abcam, ab75975) was incubated with the soluble chromatin fraction for 2 hours at 4°C. 40 μL protein G dynabeads (Invitrogen, 10003D) were then added overnight. Next, beads were sequentially washed by high-salt buffer (500 mM NaCl; 0.1% SDS; 1% Triton X-100; 2 mM EDTA; 20 mM Tris-HCl, pH 8.0) for twice, LiCl buffer (0.25 M LiCl; 1% NP-40; 10 mM Tris-HCl; pH 8.0, 1 mM EDTA) for once, and TE buffer for three times. Chromatin bound on beads was eluted twice with elution buffer (20 mM Tris-HCl, pH 8.0; 10 mM EDTA; 1% SDS) at 65 °C, each for 15 min. Eluate fraction was treated with RNase A (Thermo Fisher, EN0531), Proteinase K (Invitrogen, 25530015), and de-crosslinked at 65 °C overnight. Purified DNA was end-repaired, tailed with poly-dC, tagged with biotin, captured by Streptavidin C1 beads (Invitrogen, 65002), and ligated with an adapter. The beads-bound DNA was thoroughly washed, tagged with Illumina sequences, and sequenced with Hi-seq (2×150 bp).

Sequencing data for MCM5, CTCF, and RAD21 were processed and analyzed as reported (Liu et al., 2021b). Specifically, for MCM5, the ChIP-seq signals were normalized to fold change defined by the ratio of ChIP-ed signals over input signals. Regarding the distribution of MCM within loop domains (Rao et al., 2014), the width of loop domains was divided into 250 bins, with 20kb upstream and downstream. Matrix was generated by using normalizeToMatrix (extend=2000, w=1000, mean_mode= “absolute”, smooth=TRUE, background=0, target_ratio=25/29). Of note, a z-score-transformed fold change value was calculated for each loop domain. After that, the average z-score-transformed values at each window were used to plot the relative signal intensity at this location in the absence or presence of RAD21. For note, in RAD21 ChIP-seq, the ChIP-seq signals were normalized to the total mapped reads without removing duplications with RPKM normalization as the RAD21-depletion cells had much higher duplication rate. The parameter was used for peak calling in RAD21 ChIP-seq data: “*macs2 bdgpeakcall -c 30*”.

### PEM-seq

Cells were treated with dox for 24 hours and IAA for 6 hours before being transfected with CRISPR-Cas9 targeting the *c-MYC* or *TP53* loci., Transfected cells with or without RAD21 depletion were sorted by FACS at 48 hours post-transfection. The library preparation process is the same as previously reported (Yin et al., 2019).

PEM-seq data was processed by the PEM-Q pipeline for translocation identification as previously described (Liu et al., 2021a). Translocation was defined as genome-wide junctions excluding the off-target sites and adjacent upstream and downstream 500-kb regions. Three repeats of each treatment were combined, and translocations at the cut site chromosome were removed before SICER analysis to avoid the dominance of the cut site chromosome in cluster identification. Then, the spatial clustering approach for the identification of chromatin immunoprecipitation (ChIP)-enriched regions (SICER) algorithm (Zang et al., 2009) was applied to identify the translocation clusters with following parameters: *SICER.sh species-hg19; redundancy threshold-5; window-30,000 bp; fragment size-1; effective genome fraction- 0.74; gap size- 90, 000 bp; FDR- 0.01*. Only the clusters shared by two clones were considered as hotspot clusters. The cluster-containing genes with junctions in all 3 replicates of each sample were defined as hotspot genes.

### Quantitative 3C sequencing (q3C-seq)

Cells were treated as described in the ChIP-seq section. Briefly, G1-arrested cells with or without RAD21 were purified by FACS isolation and then fixed by formaldehyde. Fixed cells were then subjected to q3C-seq, similar to the description in 3C-HTGTS (Liu et al., 2021a), except that the 80 cycles LAM PCR were replaced by one-round primer extension and biotinylated PCR products were ligated with a barcoded bridge adapter, to quantify chromatin interaction. The 4-bp cutter DpnII (NEB, R0543L) was used for the identification of interaction from CBE upstream of *c-MYC*. The library construction procedure is similar to PEM-seq. Data of q3C-seq was applied to the PEM-Q pipeline for further processing.

### NAIL-seq

NAIL-seq was carried out as previously reported (Liu et al., 2021b) with minor modifications. For EdU -and- BrdU labeling, G1-arrested WT or RAD21-depleted cells were released into fresh medium for 3 or 4 hours, respectively. Then, cells were labeled by 20 μM EdU (Invitrogen, A10044) for 15 min, washed with pre-warmed fresh medium, and then incubated with 50 μM BrdU (Invitrogen, B23151) for 15 min. Of note, dox and IAA persisted in the medium during release and labeling with nucleoside analogs. After that, cells were thoroughly washed by ice-cold PBS and subjected to FACS purification of RAD21- depletion cells. Genomic DNA was isolated, tagged with biotin by click-iT reaction, subjected to the sequential isolation of BrdU and then EdU. After on-beads ligation, purified DNA was tagged with Illumina sequencing primers and sequenced on Hi-seq, 2×150 bp.

Data processing: NAIL-seq data were processed and early replication initiations zones (ERIZs) were identified as previously described with modifications (Liu et al., 2021b). For EdU/HU, peaks were called by MACS (version 2.1.1) with bdgpeakcall (-c 1.1 –l 200 –g 30). The neighboring peaks with an interval shorter than 10kb were merged and merged peaks with a width less than 10kb were discarded. For EdU-and-BrdU, EdU-rich regions were defined exactly the same as that in the previous report (Liu et al., 2021b). Moreover, ERIZs were also defined as EdU/HU peaks that overlapped with EdU-rich regions. Unique ERIZs in WT+IAA or RAD21-mAC+IAA were grouped into “new” or “disappeared” ERIZs after RAD21 depletion, respectively. For the common peaks, the signal intensity of EdU/HU in each peak was compared in the absence and presence of RAD21. The “increased” or “decreased” ERIZs in RAD21-depleted cells showed a 0.25-fold higher or lower EdU/HU signal intensity than that in WT cells, respectively, and the rest ERIZs were grouped into “equal”.

To depict the pattern of replication initiation for each class of ERIZs, the ERIZs were firstly centered at the midpoint with ± 100-kb regions. Then we generated the matrix using *normalizeToMatrix extend=100000, w=1000, mean_mode=“absolute”, smooth=TRUE, background=0*.

### Replication signal profile in loop domains

The contact domains in K562 WT cells were annotated in the previous study(Rao et al., 2014). Also, the loop domains were identified by the screen for domains containing the peak pixels within 50kb or 0.2 of the domain lengths at the corner (Rao et al., 2014). Non-loop domains pointed to the domains that have no overlapping regions with loop domains. Replication signals within the genomic windows of 4 classes of ERIZs were extracted and applied for metaplot analysis. All domains were divided into 250 bins with 20-kb regions upstream and downstream of both domain boundaries. The matrixes were obtained using the *normalizeToMatrix* function of R package “*EnrichedHeatmap*” with following parameters: *extend=2000, w=1000, mean_mode=“absolute”, smooth=TRUE, background=0, target_ratio=25/29.* the mean values of normalized RPKM signal were displayed for visualization.

For the computation of the distance between ERIZs and loop boundaries, the ERIZs inside loops were selected by *bedtools* for calculating the genome coordinate of their centers to the centers of closest loop boundaries. The Mann-Whitney U test was applied to compare the median value of distances. The loop boundaries were defined as the pair of loci corresponding to 10-kb pixels with high contact frequency as described (Rao et al., 2014).

K562 loop data was from GSE63525 (Rao et al., 2014).

### Repli-seq

The WT and RAD21-mAC cells were treated with dox for 24 hours, IAA for 6 or 24 hours, Hochest 33342 for 30 min, and 20 μM EdU for 15 min before harvest. The harvested cells with or without RAD21 depletion were sorted into six fractions based on the DNA content by FACS. The isolated DNA was prepared to be library following the procedure described in NAIL-seq and then sequenced on Hi-seq, 2×150 bp.

#### Data processing

Reads were processed by the NAIL-seq “RepFind” pipeline and analyzed as previously reported (Liu et al., 2021b). Briefly, reads with unique alignment, out of the blacklist regions, were normalized to RPKM signals with a 50-kb bin. The S50 value of replication timing profiles was calculated at each 50-kb bin, according to a published pipeline (https://github.com/CL-CHEN-Lab/RepliSeq), using linear interpolation of RPKM signal from G1 to G2 phase with a scale number at 100 (Brison et al., 2019).

To determine the replication timing changed regions in RAD21-depletion cells, the differences of S50 values at each 50kb bin from IAA-treated WT repeat 1 and 2 (*ΔS50wt*) were calculated. Then the differences of S50 values (*ΔS50*) between RAD21-depletion cells and WT cells were computed. The replication timing changed regions were defined as the regions in which |*ΔS50*| beyond the 95% quantile of |*ΔS50wt|*. The replication timing unchanged regions were defined as regions within the 95% quantile interval of |*ΔS50wt|*.

### PRO-seq

Cells were treated and sorted as described in the ChIP-seq section, and then subjected to nuclear run-on following the procedures in the previous report(Mahat et al., 2016). Briefly, 5 million cells, spiked with 0.25 million fruit-fly S2 cells, were incubated with ice-cold permeabilization buffer (10 mM Tris-HCl, pH 7.4, 300 mM sucrose, 10 mM KCl, 5 mM MgCl2, 1 mM EGTA, 0.05% Tween-20, 0.1% NP40, 0.5 mM DTT) supplied with PIs. The isolated nuclei resuspended in storage buffer (10 mM Tris-HCl, pH 8.0, 25% (vol/vol) glycerol, 5 mM MgCl_2_, 0.1 mM EDTA and 5 mM DTT) were subjected to nuclear run-on with biotin-11-dCTP (Jena Bioscience, NU-809-BIOX) at 37°C for 5 min. Extracted RNA was fragmented by 1 N NaOH on ice for 10 min, neutralized by 1M Tris-HCl, pH 6.8, and subjected to enrich biotinylated RNA. The nascent RNA was reversely transcribed to cDNA with an N9 random primer. After that, purified cDNA was tailed with poly-dC, biotinylated, ligated with an adapter, tagged with Illumina sequences, and sequenced on Hi-seq, 2×150 bp.

#### Data processing

The adapter sequences on R1 and R2 were trimmed by using cutadapt (-a AAGATCGGAAGAGCACACGTCTGAACTCCAGT –A NCCCCCCCCCAGATCGGAAGAGCGTCGTGTAGGGAAAGAGTGT -g GGGGGGGGGN –q 15.15 --overlap 1 –m 25). Next, reads were aligned to a combined genome (assembly hg19 combined with dm6) using bowtie (-q –very-sensitive-local –L 30 – score-min G,20,8 –reorder –rg). Duplicates or reads with multiple alignments, and reads in the blacklist regions were removed. Reads uniquely aligned to hg19 were normalized by the spike-in reads with a normalization factor measured by 1000000 / (the number of reads aligned to the dm6 genome).

Differentially expressed genes: Genes, from GENCODE, with a length > 1kb (Sigova et al., 2015) were used to quantify the transcription level by using featureCounts (-p –M –F SAF –s 2). The differential expression testing was performed using DESeq2 bioconductor package with a threshold of Benjamini-Hochberg corrected *p*-value < 0.01, and log2 (FC) > 1.

## Data availability

All sequencing data presented in the present study were deposited at NCBI Gene Expression Omnibus (GEO) under the accession number: GSE189762.

## Contributions

J.H. conceived and supervised the project. J.W., Y.L., X.L., and J.H. designed the experiments; J.W., Y.L., X.L., T.G., Y.L., J.Y., and W.Z. performed the experiments; J.W., Y.L., Z.Z., X.L., C.A., and J.H. analyzed the data; J.W., Y.L., Z.Z., and J.H. wrote the paper.

**Table 1.**
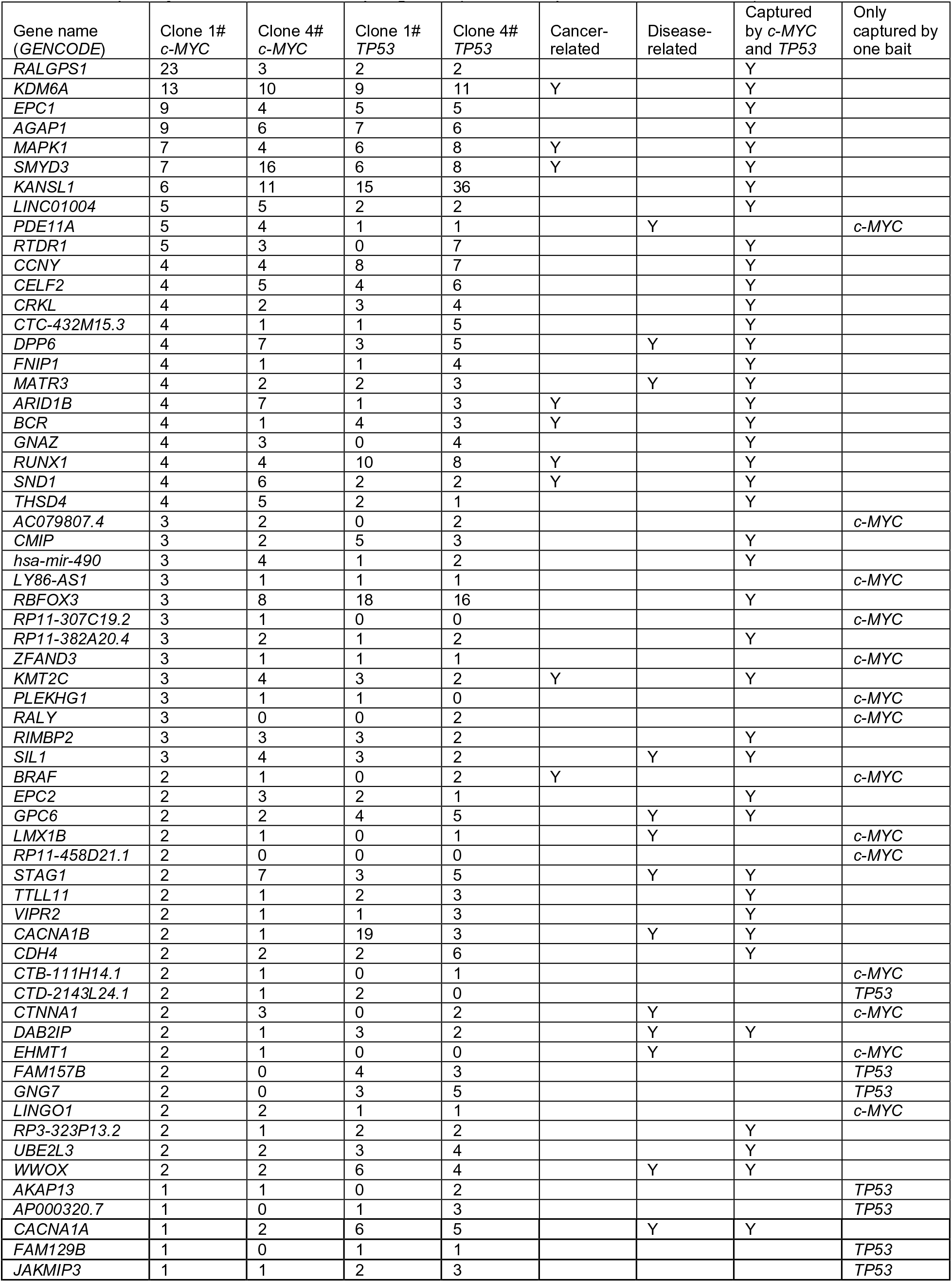
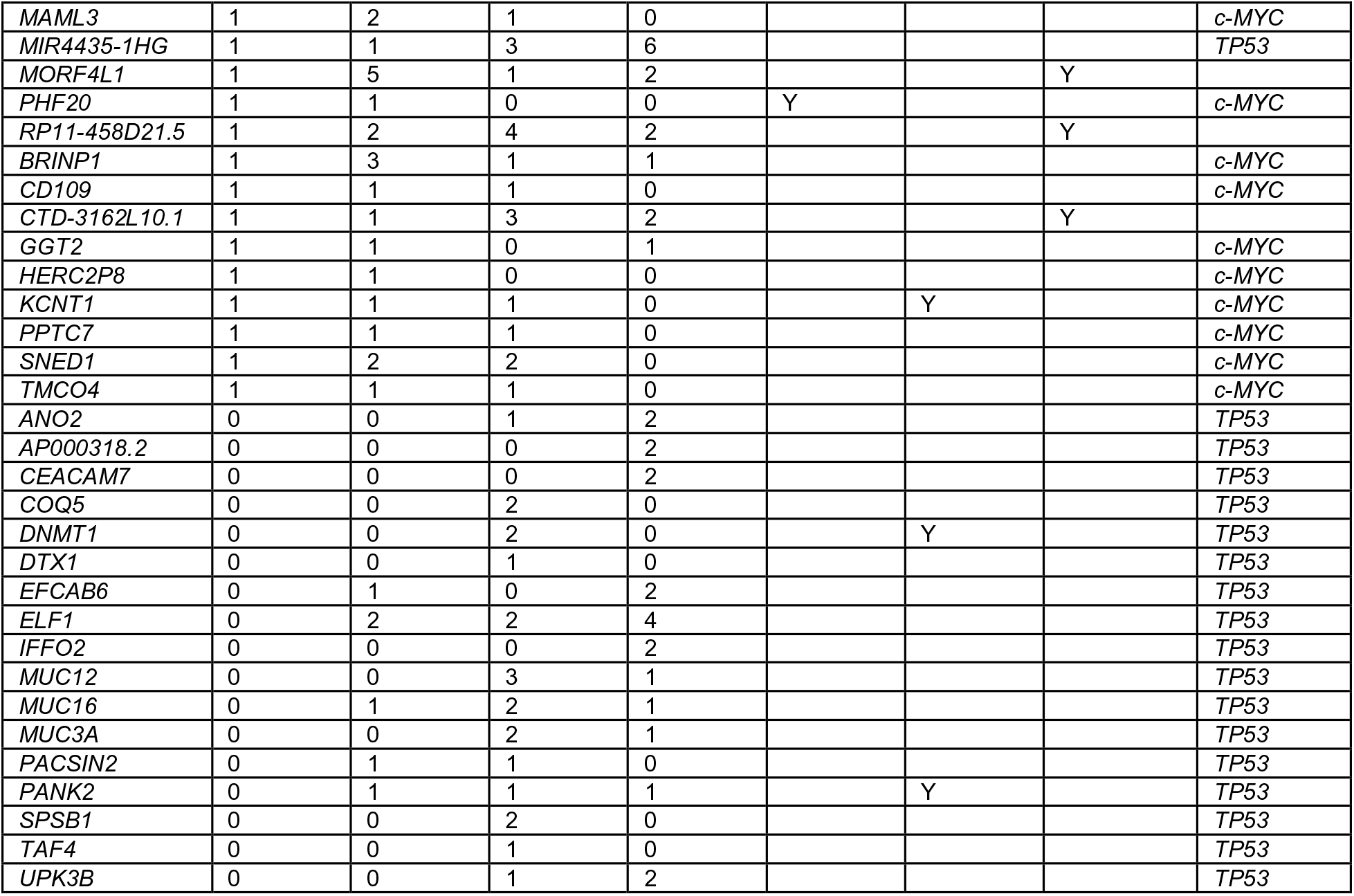
Frequency of translocation hotspot genes. summarized the translocation frequency of hotspot genes captured by c-MYC and TP53 DSBs. The bait loci and genes are indicated in italic font. The genes that have 3 or more junctions captured by both baits were noted in the 8^th^ column. Y, yes.

## Acknowledgments

We acknowledge the members of the Hu laboratory for their helpful discussions. This work was supported by the National Key R&D Program of China (2017YFA0506700 to J.H.) and the NSFC grant (31771485 and 32122018 to J.H.). J.H. is an investigator at the PKU-TSU Center for Life Sciences. Y.L. is supported by the Boehringer Ingelheim-Peking University Postdoctoral Program. We thank the Flow Cytometry Core at the National Center for Protein Sciences, Peking University. The degron plasmid backbones were kind gifts from Dr. Xiong Ji.

## Supplemental Figure Legends

**Figure S1.**
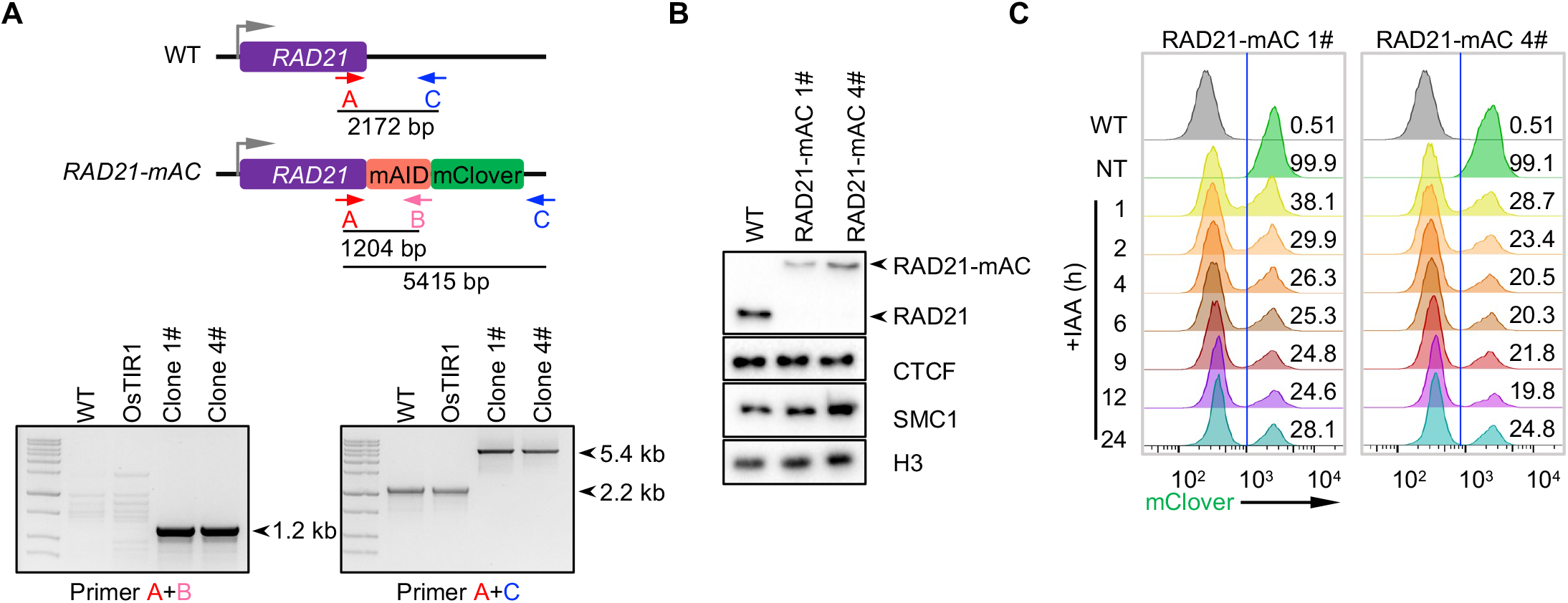
**Characterization of RAD21-mAC K562 cells** A. Knock-in and PCR screening strategy for the RAD21 tagged with mAID-mClover. The result of PCR validation was showed on the bottom. The amplified products were confirmed by sanger sequencing. B. Western blotting for chromatin-bound RAD21, CTCF and SMC1 using histone H3 as loading control in WT and RAD21-mAC cells (no IAA treatment). C. Flow cytometry analyzing mClover fluorescence of two RAD21-mAC clones after IAA treatment for indicated time.

**Figure S2.**
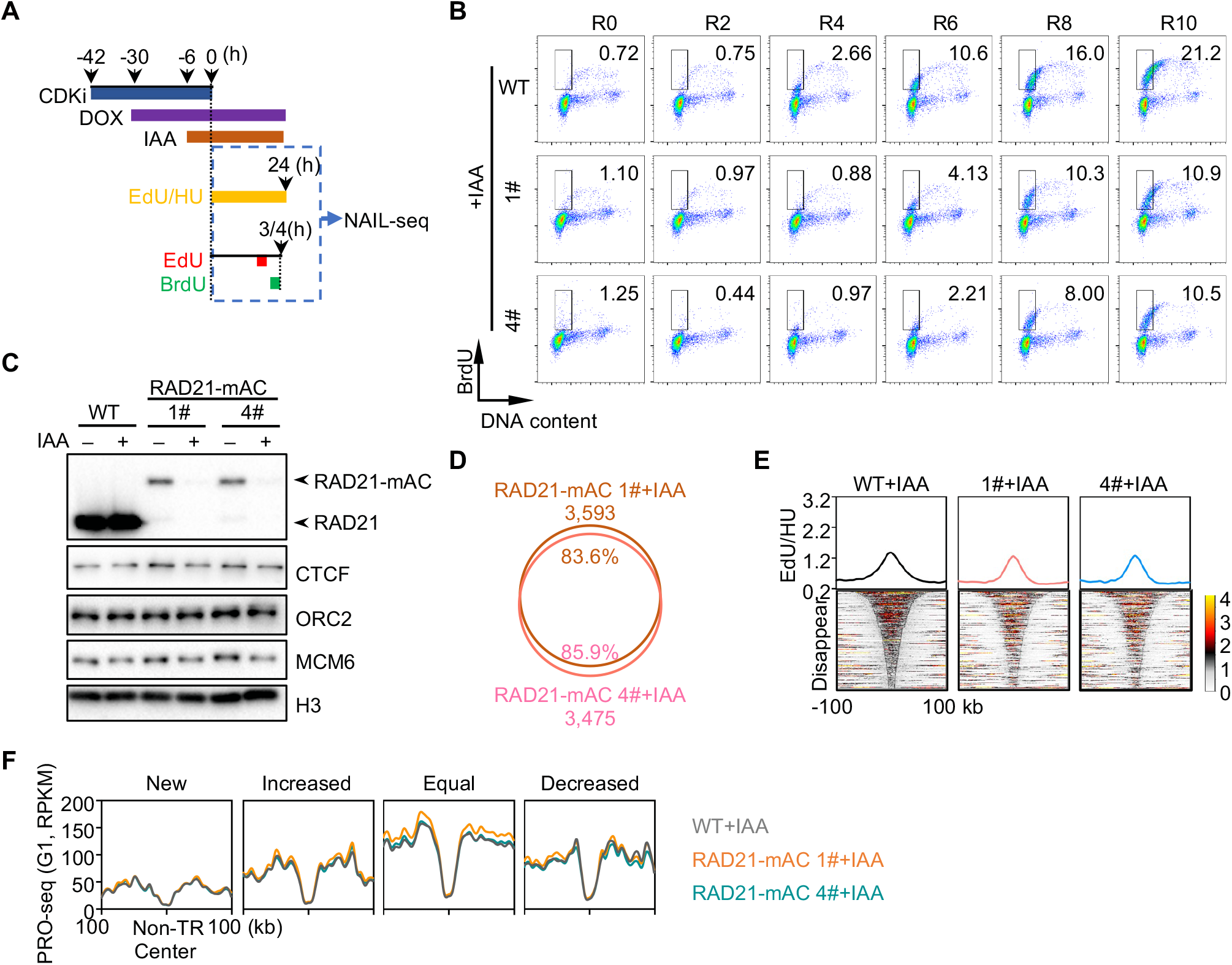
**RAD21 regulates the firing of early replication origins in K562 cells** A. Schematic for NAIL-seq analysis to identify early replication initiation zones (ERIZs). The detailed procedures are described in Methods. B. The release of G1-arrested WT and RAD21-depleted into early S phase. Cells were synchronized by CDKi for 36 hours, treated with IAA for 6 hours before being released into a fresh medium with dox and IAA. Cells were labeled with BrdU for 30 minutes before being harvested at the indicated time and then fixed by ethanol. Rx, Release for x hours. C. Western blotting showing the amounts of chromatin-bound RAD21, CTCF, ORC2, and MCM6 in the G1-arrested WT or RAD21-mAC cells with IAA treatment for 6 hours. D. Overlap of ERIZs identified in IAA-treated RAD21-mAC clones 1# and 4#. E. Heatmap showing the intensity of early replication signals from IAA-treated WT and RAD21-mAC cells in disappeared ERIZs. F. Signal intensity of PRO-seq from G1-arrested cells with or without RAD21 depletion in non-transcribed regions adjacent to each class of ERIZs.

**Figure S3.**
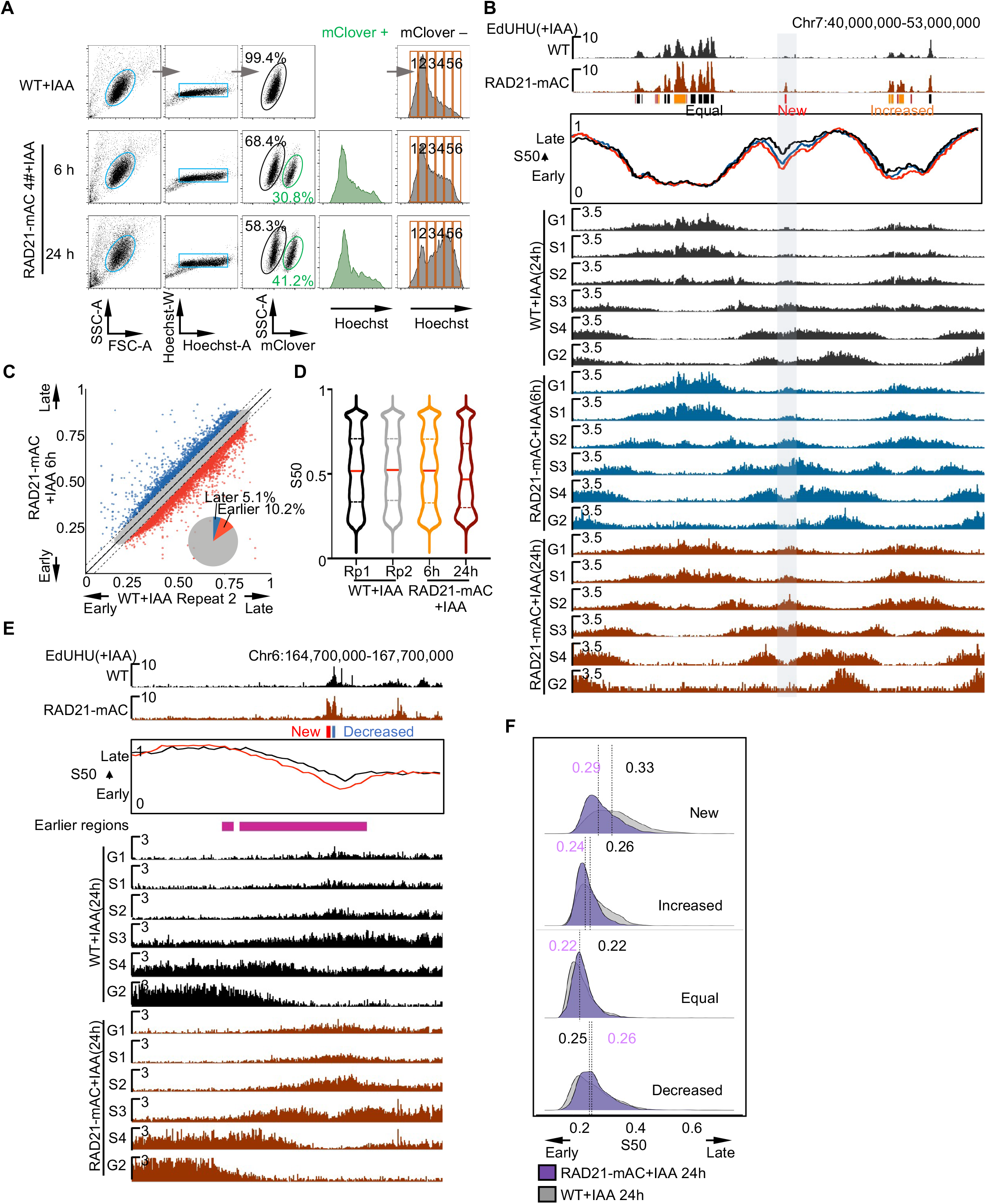
**RAD21 depletion affects replication timing in K562 cells.** A. FACS sorting strategy for replication timing analysis. mClover negative live cells from IAA treated WT and RAD21-mAC cells were gated by blue boxes, the black and green circles gate the mClover negative and positive cells, respectively. The red rectangles indicate the 6 fractions sorted according to DNA content of cells marked by Hoechest33342. B. A representative profile of replication timing in IAA-treated WT and RAD21-mAC cells (clone 4#). Grey shadow highlights the neighbor region of a new ERIZ. C. Scatter plot of S50 values between WT and RAD21-mAC cells (clone 4#) with IAA treatment for 6 hours. Legends are described in Figure 3C. D. Distribution of S50 values in IAA-treated WT and RAD21-mAC cells. Rp, repeat. E. A representative profile of a decreased ERIZ overlapping with a region showing an earlier replication timing. F. Density plots showing the distribution of S50 values from IAA-treated WT (grey) or RAD21-mAC (purple) cells in each class of ERIZs identified in RAD21-mAC 4#. The dashed lines and number mark the median of S50 in WT (grey numbers) or RAD21-mAC (purple numbers) cells.

**Figure S4.**
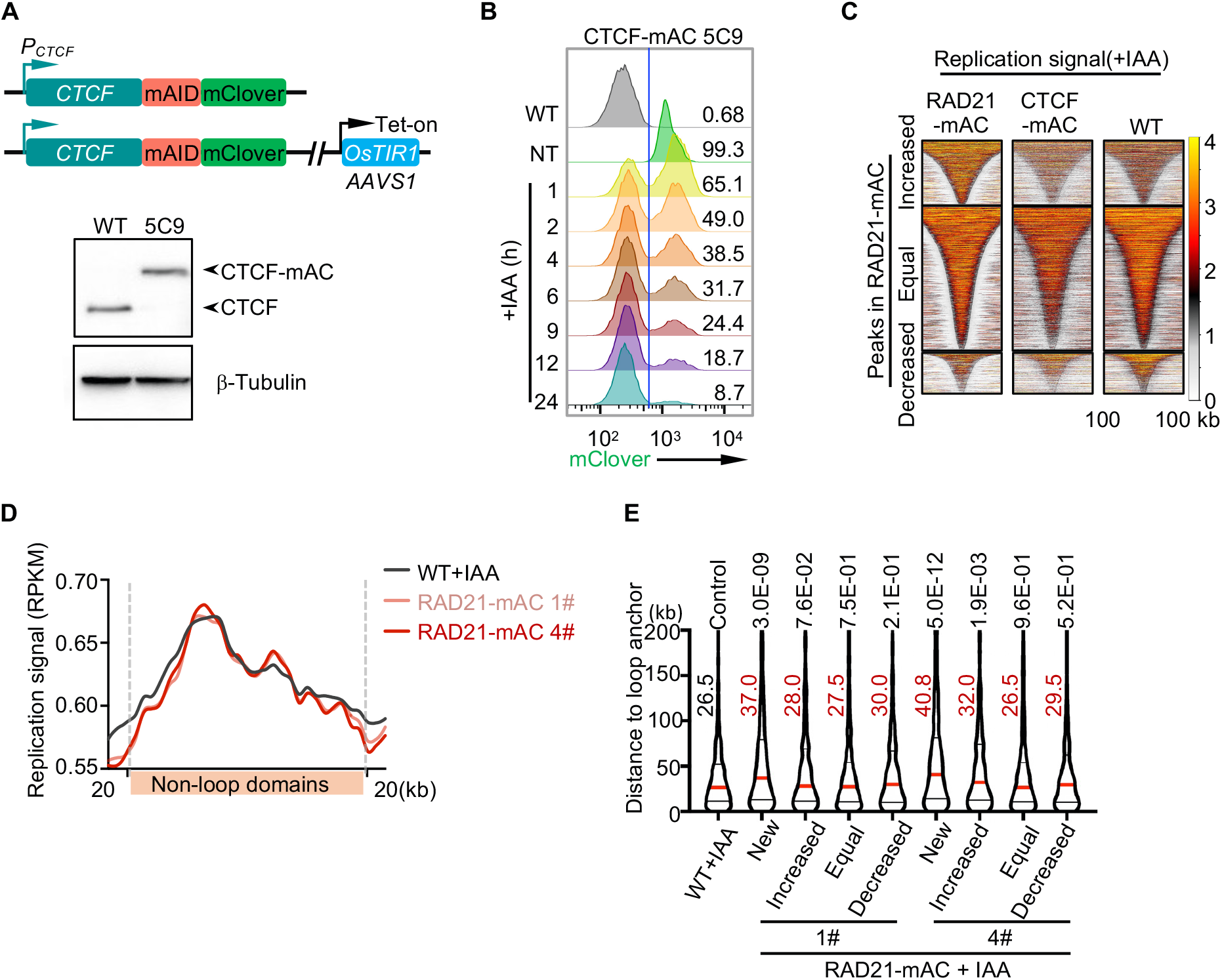
**The distribution of early replication initiation sites after RAD21 depletion** A. Western blotting for the validation of the CTCF-mAC cell line (clone 5C9). B. Flow cytometry analyzing mClover fluorescence of CTCF-mAC clone 5C9 after IAA treatment for indicated time. C. Heatmaps showing the distribution of early replication signals from IAA-treated RAD21-mAC, CTCF-mAC, and WT samples at the increased, equal, and decreased ERIZs of RAD21-mAC 1#. The ERIZs were ranked by width and centered at the midpoint. Legends are depicted as described in Figure 2D. D. Distribution of early replication signals in the presence (black) or absence (pink or red) of RAD21 within non-loop domains of WT cells. Each loop domain was divided into 250 bins and aligned at loop boundaries, with 20 kb regions upstream and downstream. E. Distances between the center of each ERIZ and the closest loop anchor. ERIZs outside of loops were excluded for analysis. Red lines and numbers mark the median distances. The *p*-value from Mann–Whitney U test is labeled on the top of the violin plots.

**Figure S5.**
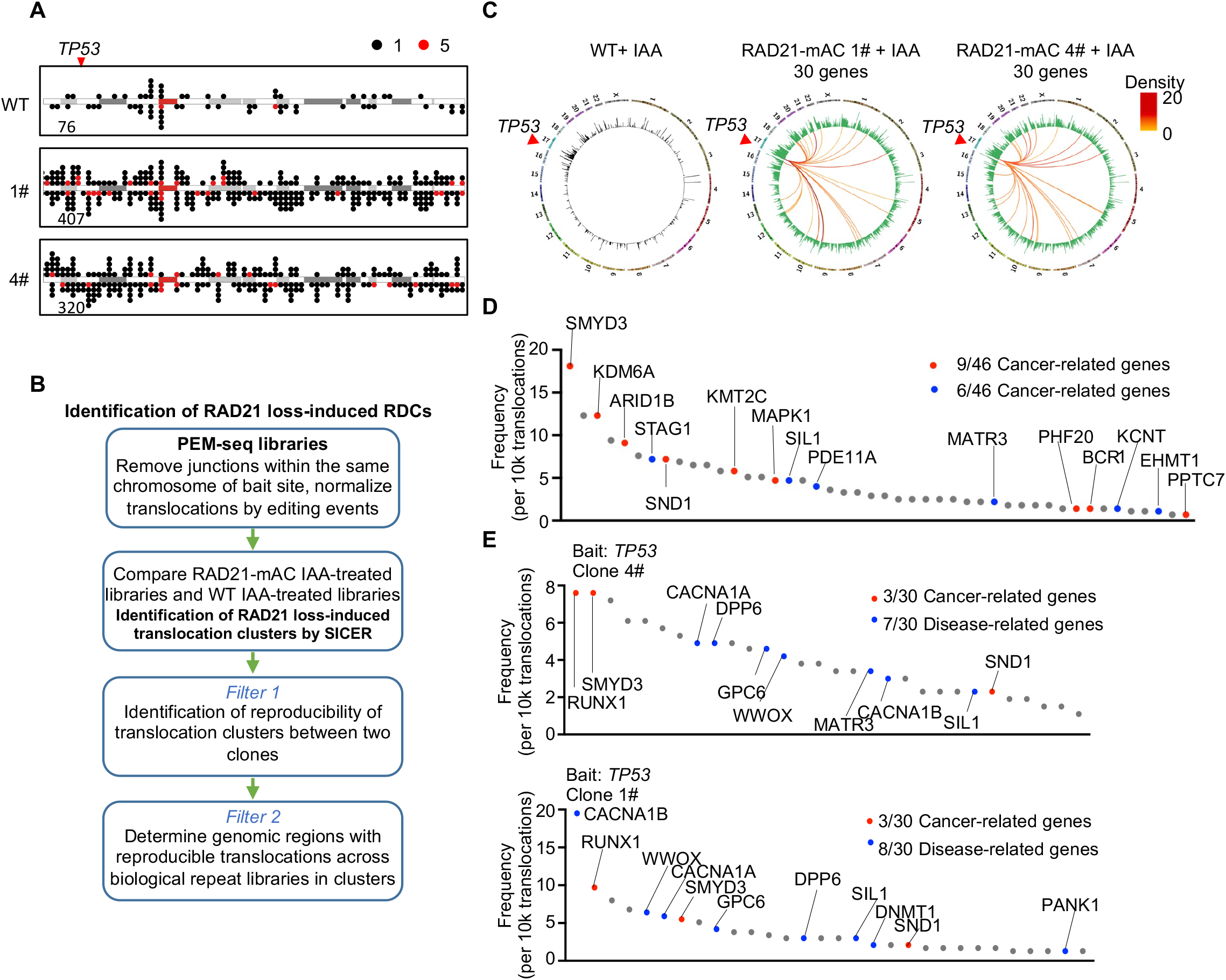
**RAD21 depletion leads to genome instability** A. Dot plots exhibiting the distribution of translocation junctions on Chromosome 17 in IAA-treated cells. The number of translocations were normalized to 100,000 editing events from each sample. The red arrow indicates the position of bait site at the *TP53* locus. Numbers of junctions are indicated at each plot. Legend for dot colors is indicated on the top. B. Flow diagram for the identification of translocation hotspot genes. See Methods for more details. C. Circos plots showing the genome-wide translocations and translocation hotspot genes cloned from the *TP53* locus with RAD21 depletion. The legend of the circos plot is described in Figure 5D. D. Translocation frequencies of RAD21-depletion-induced translocation hotspot genes in RAD21-mAC 4# captured by *c-MYC* DSBs. Legends are depicted as described in Figure 5E. E. Translocation frequencies of RAD21-depletion-induced translocation hotspot genes in RAD21-mAC 1# (top) and 4# (down) captured by *TP53* DSBs.

**Figure S6.**
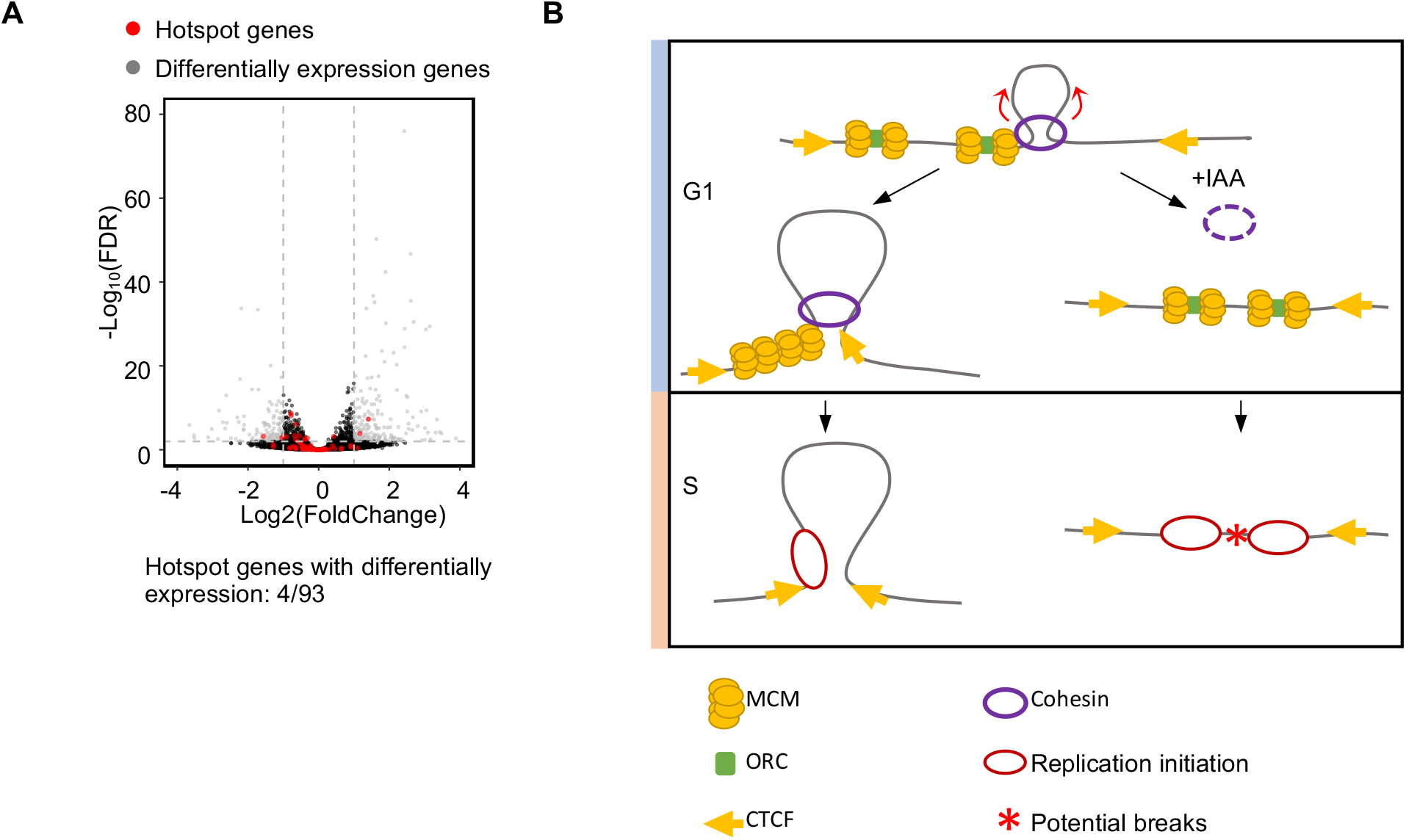
**Translocation hotspot genes tend to locate in replication timing-altered regions** A. Volcano plot depicting the fold change of gene expression in the absence or presence of RAD21. Red dots mark the translocation hotspot genes identified by *c-MYC* and *TP53* DSBs; grey dots mark the differentially expressed genes after RAD21 depletion in the G1 phase. B. Working model of cohesin-mediated loop extrusion in modulating early replication initiation. During the G1 phase, chromatin-bound MCM double hexamer undergoes redistribution mediated by cohesin-driven loop extrusion and stalls at loop boundaries, resulting in early replication adjacent to the loop boundaries in WT cells. Loss of cohesin-mediated MCM redistribution induces ectopic origin firing within loop domains but away from loop boundaries in the early S phase. The firing of multiple origins in a limited regions may lead to DSBs as indicated by red asterisk.

## Key resource table

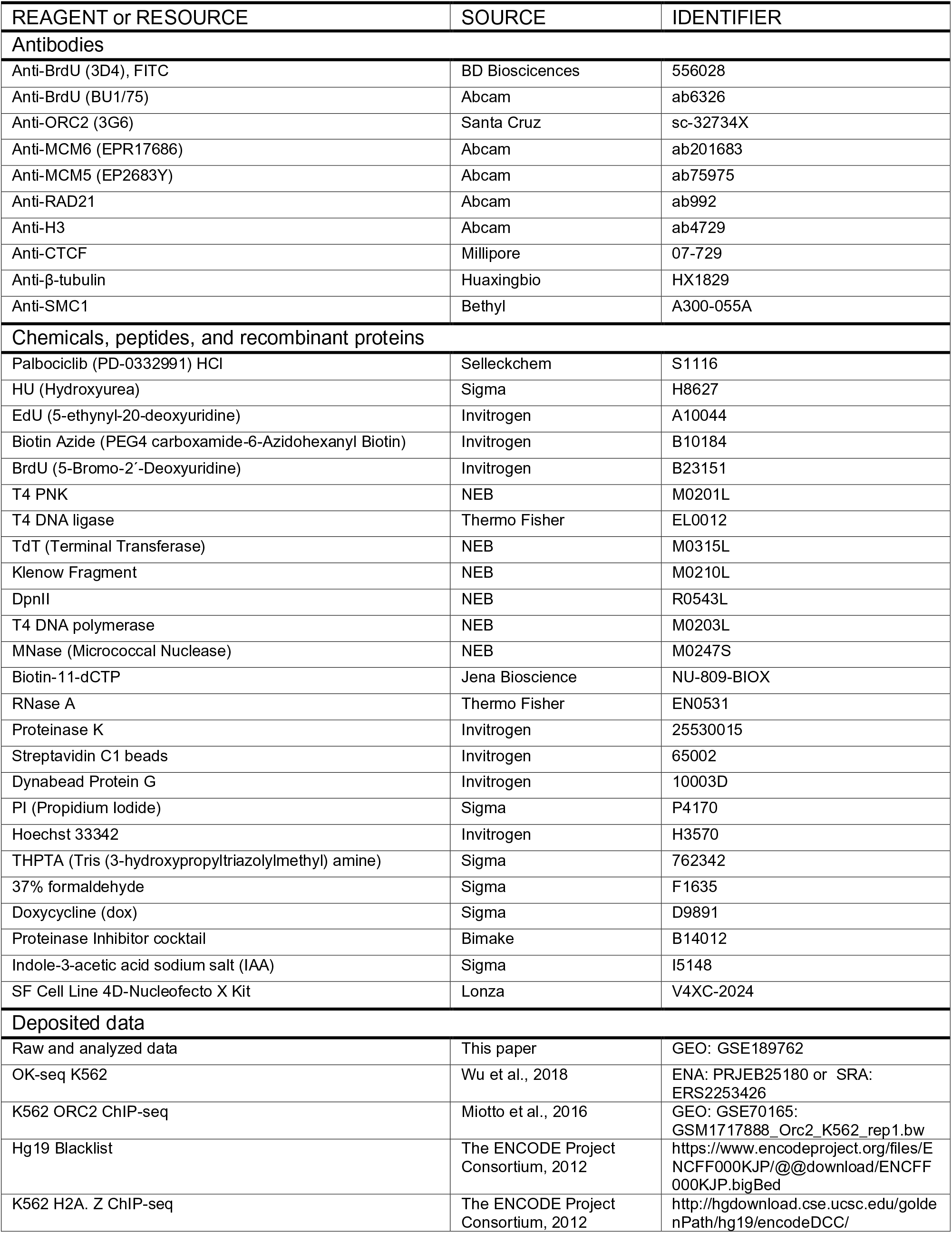
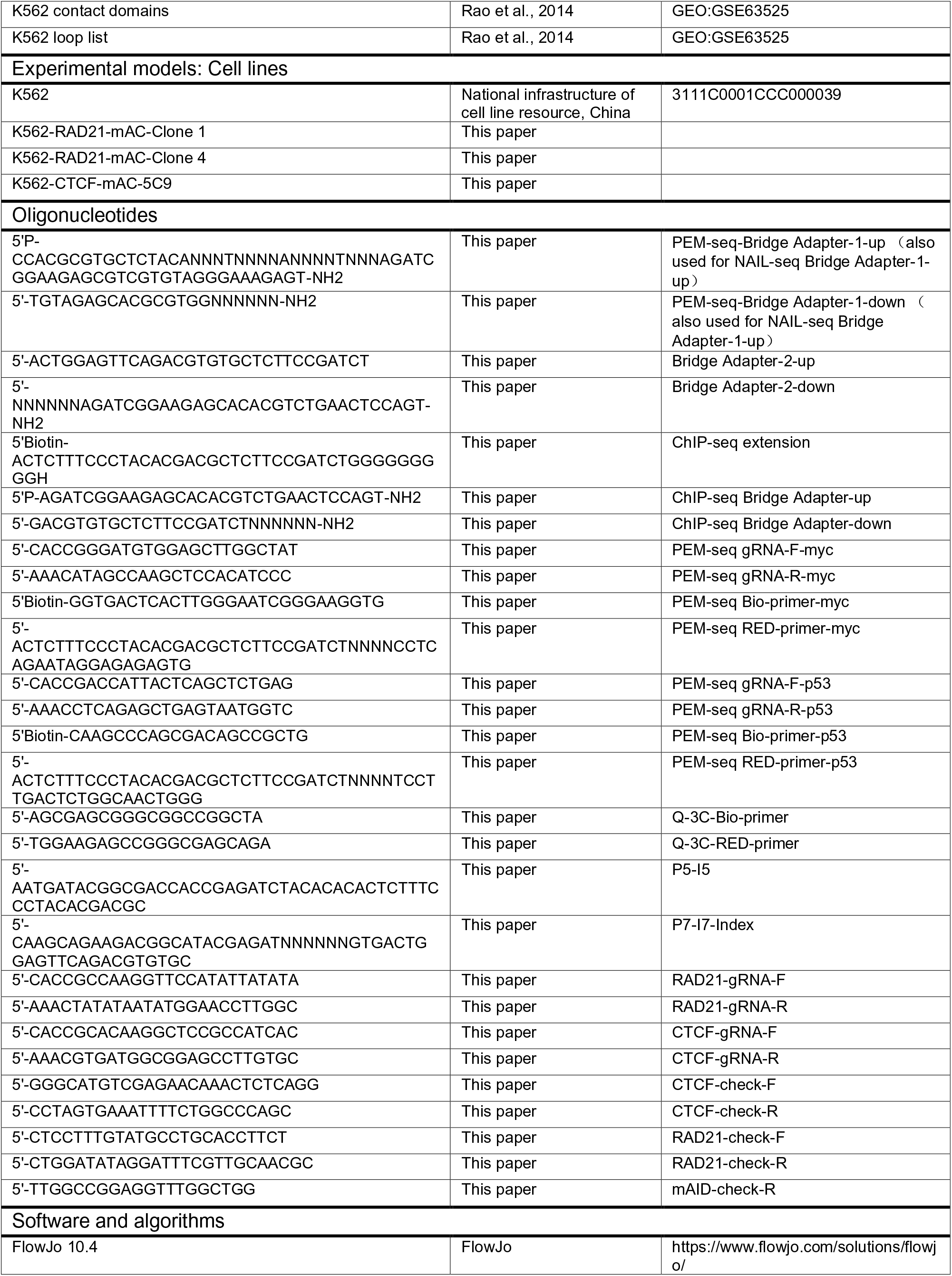
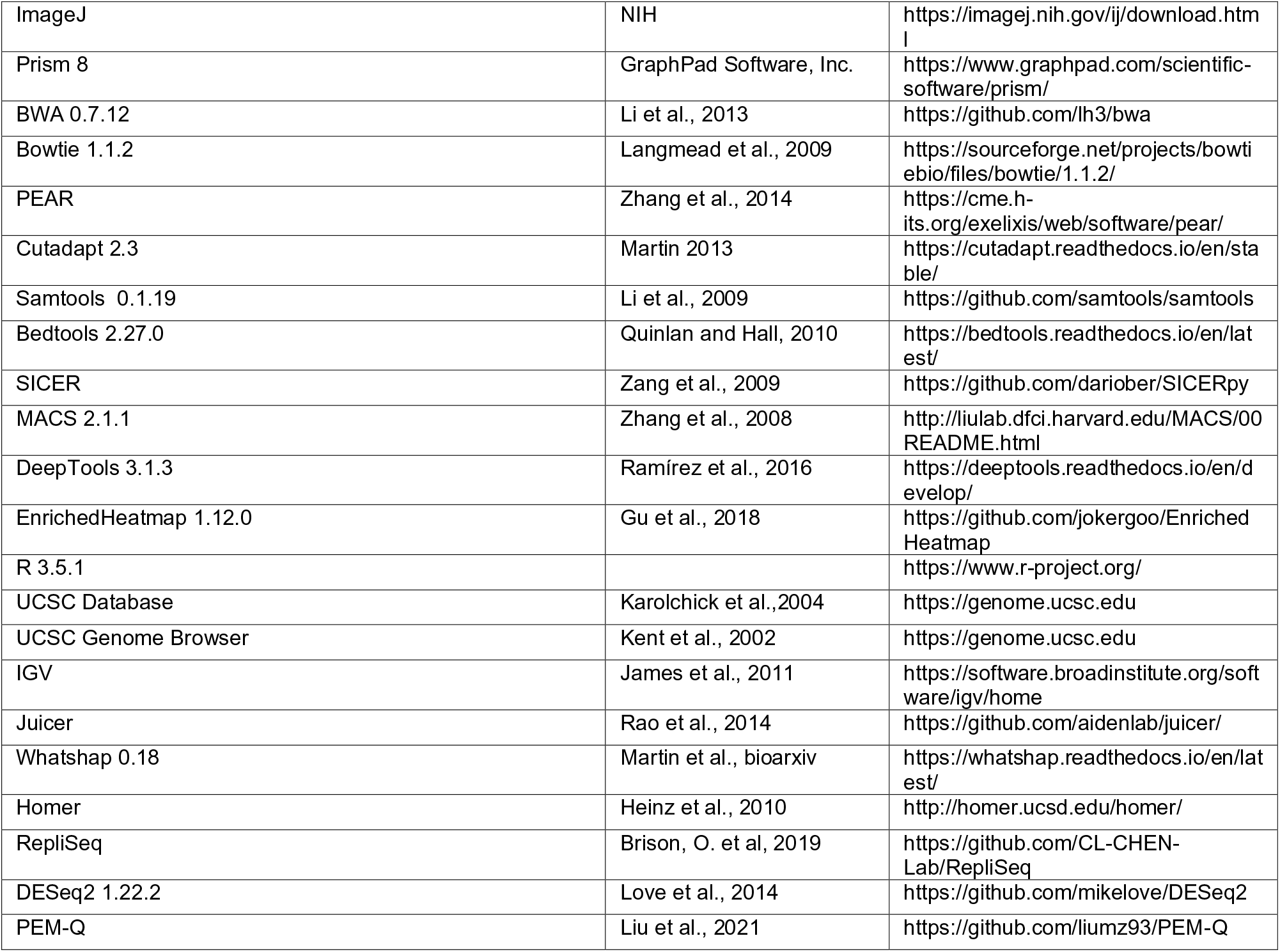
XXX

